# Cannabinoid receptor 1 activity promotes microbial dysbiosis by stimulating the oxygen-dependent expansion of Enterobacteriaceae

**DOI:** 10.64898/2026.10.10.758071

**Authors:** Mary K. Mitchell, Mikayla A. Carty, Nayeli L. Snow, Mary T. Archambault, Addison E. Budsberg, Elisa V. Carrillo, Emily E. Vignieri, Ethan A. Older, Olivia L. Rometo, Lauren E. Van de Wal, Kayla J. Floyd, Lindsey R. Baker, Claire A. Kunkle, Jie Li, Melissa Ellermann

## Abstract

The endocannabinoid system is comprised of bioactive lipids that signal via cannabinoid receptors, which can modulate barrier and immune functions along the gastrointestinal tract. Clinical studies indicate that endocannabinoid activity is dysregulated in inflammatory bowel diseases. Moreover, an increase in gut endocannabinoids is correlated with microbial dysbiosis in Crohn’s disease. Here, we show that endocannabinoid activity promotes metabolic dysfunction within the colonic mucosa, resulting in microbial dysbiosis and exacerbated colitis. Stimulation of the cannabinoid receptor CB1 resulted in the expansion of Enterobacteriaceae within the gut microbiome, a common feature of inflammation-associated dysbiosis. Preceding this microbial change, CB1 activity induced transcriptional signatures of mitochondrial stress and immune activation within the colonic mucosa. This was followed by a reduction in mucosal ATP production and the induction of aerobic respiration genes in luminal *Escherichia coli*. Genetic inactivation of aerobic respiration in *E. coli* limited its proliferation, indicating that CB1 activity stimulates oxygen-dependent Enterobacteriaceae growth. In IBD susceptible mice, endocannabinoid activity promoted mucosal infection by an *E. coli* pathobiont, which corresponded with elevated pro-inflammatory markers. In inflammation resistant mice, acute CB1 stimulation was sufficient to accelerate the infection dynamics of an Enterobacteriaceae pathogen, resulting in exacerbated colitis. Taken together, our data support a model in which endocannabinoid signaling at the CB1 receptor disrupts respiratory metabolism in the colon, compromising the anoxic environment of the lumen that is critical for maintaining aerobically respiring pathogens at low abundance. More broadly, our findings highlight a proinflammatory effect of endocannabinoid activity that can contribute to intestinal disease development.

## Introduction

Inflammatory bowel diseases (IBD), including Crohn’s disease and ulcerative colitis, are chronic and relapsing diseases with complex etiologies that involve host genetic susceptibility, environmental factors, and microbiome dysfunction. The development of IBD is influenced by pathological responses to and insufficient healing from acute mucosal injury, which can lead to sustained immune activation and chronic tissue damage.^1^ In conjunction, pathological changes to the gut microbiome -termed dysbiosis – can further contribute to disease development. This is often characterized by a decreased capacity for establishing colonization resistance, an ecological process by which the host microbiome limits the proliferation of resident pathobionts and invading pathogens.^2^ Indeed, a common feature of IBD-associated dysbiosis includes the overgrowth of pathobionts that contribute to barrier dysfunction and excessive proinflammatory responses, which often coincides with the contraction of bacterial families that are critical for maintaining mucosal homeostasis.^3^

Metabolic processes within the colonic epithelium strongly influence the symbiotic or dysbiotic nature of the gut microbiome.^1,4^ Under homeostasis, colonocytes typically utilize butyrate as an energy source via mitochondrial beta-oxidation and oxidative phosphorylation.^5^ This oxygen consuming process induces a state of physiologic hypoxia within the colonic epithelium and helps establish the anoxic luminal environment, which favors the growth of obligate anaerobes.^6,7^ Butyrate is a microbial product that is generated via the fermentation of dietary fibers by obligate anaerobes such as Lachnospiraceae, thus establishing a positive feedback loop that helps maintain microbial symbiosis.^8^ During inflammation, reprogramming of host metabolism can disrupt these symbiotic interactions and favor the overgrowth of bacterial populations with pathogenic potential. In the inflamed colon, a glycolytic shift in colonocytes decreases oxygen consumption and enables its diffusion into the lumen, which supports the proliferation of facultative anaerobes such as Enterobacteriaceae.^6,9–12^ Subsequent inflammatory processes further remodel the nutrient landscape to sustain microbial dysbiosis. Indeed, blooms of intestinal Enterobacteriaceae represent a common feature of inflammation-associated dysbiosis reported in Crohn’s disease and in experimental models of colitis.^13–18^ Thus, the capacity of the microbiome to establish colonization resistance is substantially diminished in the inflamed gut, which can enable the exacerbation of disease by pathogenic Enterobacteriaceae following their luminal proliferation and infection of the mucosa.^19^

The endocannabinoid system is comprised of bioactive lipids known as endocannabinoids, their cognate cannabinoid receptors, and host biosynthetic and degradative enzymes that regulate local tissue concentrations of these molecules. In the gut, endocannabinoid signaling at the CB1 and CB2 cannabinoid receptors influences epithelial barrier integrity and immune functions, which can impact inflammation outcomes.^20,21^ The CB1 receptor is largely expressed within the enteric nervous system and in a subset of epithelial cells, while CB2 is primarily expressed by immune cells.^20^ In genetic models of IBD, repeated stimulation of CB2 once disease is established attenuates chronic inflammation by decreasing mucosal infiltration of proinflammatory immune cells and activation of effector T-cells while augmenting regulatory immune responses.^22,23^ Similarly, in models of chemically induced epithelial injury, CB1 and CB2 activity are protective during acute colitis development.^22,24–35^ As a result, endocannabinoid activity within the gut has been largely characterized as anti-inflammatory.

In human IBD, clinical studies have repeatedly correlated disease status with alterations in endocannabinoid tone.^23,35–41^ However, the specific effects differ among IBD subtypes and between quiescent versus active disease. In Crohn’s disease, an increase in both intestinal and circulating endocannabinoids and an elevation in CB1 expression in gut tissues have been reported.^23,36–39^ Moreover, untargeted metabolomics performed as part of the Integrative Human Microbiome Project (iHMP) have revealed that fecal arachidonoyl lipids including endocannabinoids are significantly elevated during active Crohn’s disease compared to remission and non-inflamed controls.^42–44^ This is positively correlated with established features of microbial dysbiosis in Crohn’s disease including Enterobacteriaceae overgrowth. However, it remains unclear whether this increase in endocannabinoid tone is simply an indicator of inflammation, or whether these changes signify elevated endocannabinoid activity that directly impacts Crohn’s disease pathogenesis.

Here, we show that endocannabinoid activity at the CB1 receptor provokes acute metabolic injury within the colonic mucosa, resulting in microbial dysbiosis and exacerbated colitis development. Using mouse models, we show that acute stimulation of CB1 activity in the uninflamed gut promotes the rapid expansion of resident *Escherichia coli* and other facultative anaerobes within the gut microbiome. Preceding these microbial changes, CB1 activity induced transcriptional signatures of mitochondrial stress and immune activation within the colonic mucosa. This was followed by a reduction in mucosal ATP production and the induction of aerobic respiration genes in luminal *E. coli* that coincided with dysbiosis. Genetic inactivation of aerobic respiration in *E. coli* limited its proliferation, indicating that CB1 activity stimulates oxygen-dependent *E. coli* growth. In IBD susceptible mice, CB1 activity promoted mucosal infection by adherent-invasive *E. coli* – a pathobiont associated with Crohn’s disease, which corresponded with elevated pro-inflammatory markers in the colon. In inflammation resistant mice, acute CB1 stimulation was sufficient to accelerate the infection dynamics of *Citrobacter rodentium*, a pathogenic facultative anaerobe, resulting in exacerbated colitis. Taken together, our data support a model in which endocannabinoid signaling at the CB1 receptor disrupts host respiratory metabolism within the mucosa, compromising the anoxic environment of the lumen that is critical for maintaining aerobically respiring pathogens at low abundance, resulting in microbial dysbiosis. More broadly, our findings highlight a novel proinflammatory effect of host endocannabinoid activity that can contribute to the development of intestinal disease.

## Results

### Host endocannabinoid activity induces the proliferation of endogenous *Escherichia coli* within the gut microbiome

Considering the positive correlation between fecal endocannabinoids and microbial dysbiosis reported in Crohn’s disease ^42–44^, we first investigated whether acutely increasing host endocannabinoid tone can alter microbiome composition. To accomplish this, we utilized an irreversible inhibitor that blocks the activity of monoacylglycerol lipase (Magl), a host enzyme that hydrolyzes the endocannabinoid 2-arachidonoyl glycerol (2-AG).^45^ Acute Magl inhibition promotes the accumulation of 2-AG within intestinal tissues,^32,46^ which we confirmed in our mouse model **(Fig S1A)**. We first tested the effects of Magl inhibition in non-inflamed WT mice because inflammation can modulate nutrient availability within the gut, thus altering microbiome composition.^47^ Because we utilized mice that lack endogenous Enterobacteriaceae within their microbiomes, we engrafted a gut commensal *E. coli* isolate prior to our treatments that stably colonizes the murine gut without prior antibiotic treatment **(Fig 1A, S1B)**. In contrast to vehicle controls, treatment with the Magl inhibitor increased fecal *E. coli* abundance in male and female mice by orders of magnitude within 12 hours **(Fig 1B, S1C)** and in a dose dependent manner **(Fig 1C)**. Fecal *E. coli* remained significantly elevated up to 48 hours post treatment **(Fig 1B)**. Together, our results suggest that increasing host endocannabinoid activity stimulates the transient proliferation of *E. coli* within the gut microbiome.

**Fig. 1.**
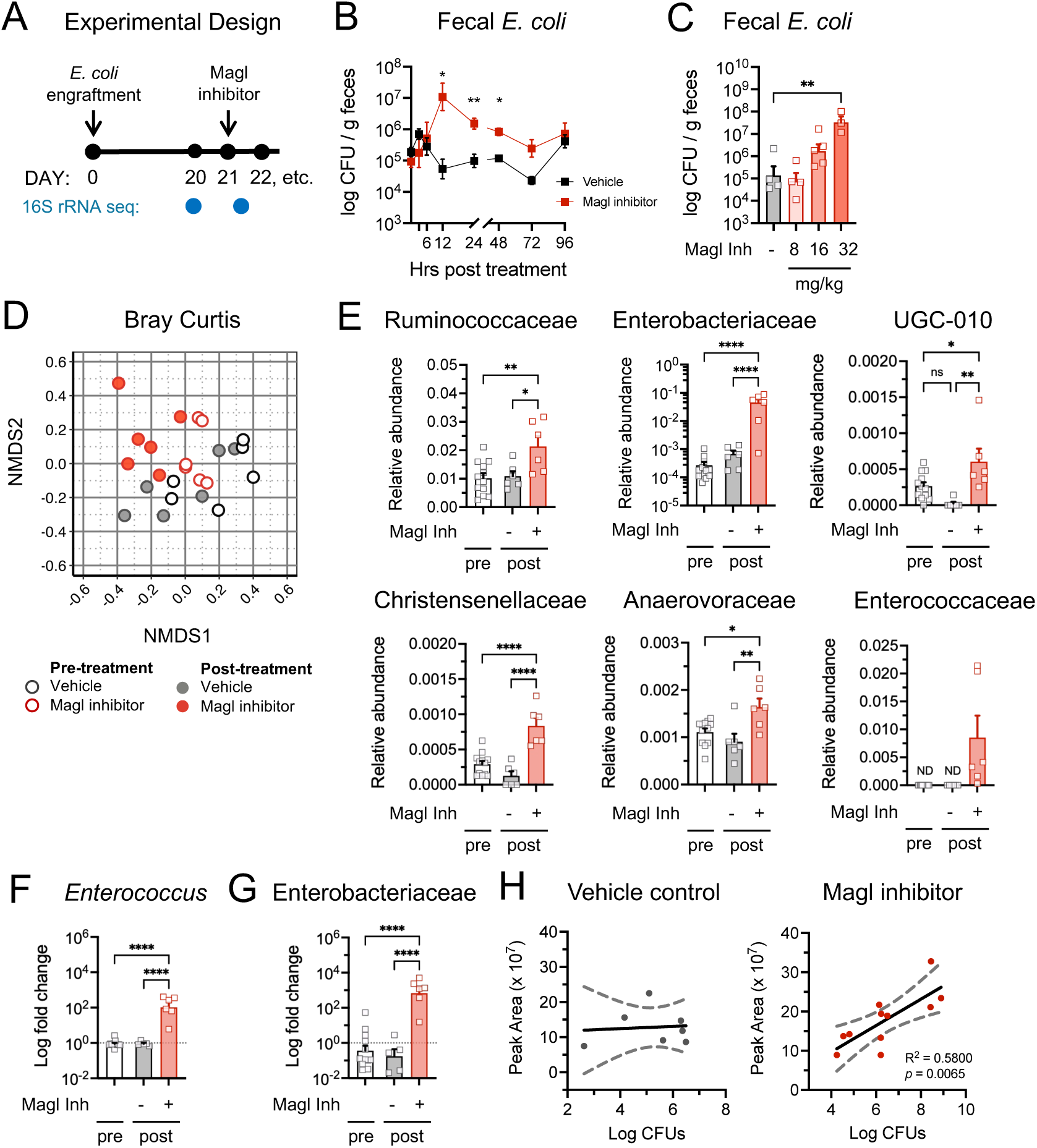
Host endocannabinoid activity induces the proliferation of endogenous *Escherichia coli* within the gut microbiome. A) Experimental design for all panels. B) Female mice were administered a single dose of vehicle or Magl inhibitor (32 mg/kg). Data depict fecal *E. coli* CFUs post-treatment (*n* = 10-11 per group, pooled from 2 cohorts). Two-way ANOVA with Sidak’s multiple comparison test. C) Female mice were administered a single dose of vehicle or Magl inhibitor. Data depict fecal *E. coli* CFUs at 24 hours post-treatment (*n* = 3-5 per group). One-way ANOVA with Tukey’s multiple comparison test. D-J) Female mice were administered a single dose of vehicle or Magl inhibitor (32 mg/kg). Fecal pellets were collected pre-treatment and 12 hours post-treatment (*n* = 6 per group distributed in 3 cages). D) Bray-Curtis dissimilarities of fecal microbiomes post-treatment. E) Relative fecal abundance of significantly altered families pre- and post-treatment. F-G) Relative fecal abundance of F) *Enterococcus* and G) Enterobacteriaceae as determined by targeted qPCR. Data are represented as log fold change relative to pre-treatment. Due to insufficient DNA, one sample was excluded from the vehicle post-treatment group. One-way ANOVA with Tukey’s multiple comparison test. ND, not detected. H) Female mice were administered a single dose of vehicle or Magl inhibitor (32 mg/kg). Detection and quantification of 2-AG by LC-MS in colon tissues, and fecal culture of *E. coli*, at 15-17 hours post-treatment (*n* = 7 or 11 per group, pooled from 2 cohorts). Simple linear regression of colon 2-AG and fecal *E. coli* abundance with vehicle treatment or Magl inhibition. * *p* < 0.05, ** *p* < 0.01, **** *p* < 0.0001.

We next sought to evaluate the effects of endocannabinoid activity on overall gut microbiota composition. 16S rRNA sequencing revealed that acute Magl inhibition significantly altered the composition of the microbiota relative to vehicle controls (PERMANOVA *F* = 2.50, *p* = 0.011) and pre-treatment controls (PERMANOVA *F* = 2.42, *p* = 0.039) (**Fig 1D**). In contrast, microbiota composition was not altered in the vehicle group relative to pre-treatment controls (PERMANOVA *F* = 1.33, *p* = 0.22). Family level analysis revealed five bacterial families in addition to Enterobacteriaceae that were significantly increased following Magl inhibition: Ruminococcaceae, UCG-010, Enterococcaceae, Christensenellaceae, and Anaerovoracaceae (**Fig 1E, Table S1**). Notably, while Enterococcaceae was below detection in pre-treatment and vehicle samples by 16S rRNA sequencing, Magl inhibition increased its relative abundance above the detection limit in 5 out of 6 mice (**Fig 1E**). Using a more sensitive approach, targeted qPCR confirmed the relative expansion of *Enterococcus* and Enterobacteriaceae in feces following Magl inhibition (**Fig 1F-G**). Selective culture further confirmed that the absolute abundance of *E. coli* in cecal contents and feces also increased in these mice by several orders of magnitude (**Fig S1D**). In contrast, we did not detect any bacterial families that significantly contracted following Magl inhibition (**Table S1**). This corresponded with a significant increase in alpha diversity in the Magl inhibitor group compared to pre-treatment controls (**Fig S1E**). Considering the positive association between endocannabinoids and gut Enterobacteriaceae reported in Crohn’s disease ^42–44^, we lastly determined whether the extent of *E. coli* proliferation could be correlated with colon 2-AG concentrations in our model. In mice treated with the Magl inhibitor, fecal *E. coli* abundance positively correlated with 2-AG levels in colon tissues **(Fig 1H)**. In contrast, no correlation was observed between basal colon 2-AG and fecal *E. coli* in vehicle controls. Together, these observations demonstrate that increasing endocannabinoid activity alters the gut microbiome by promoting the marked expansion of select bacterial taxa usually present at low abundance. This includes two families of facultative anaerobes – Enterobacteriaceae and Enterococcaceae – that encompass inflammation-associated pathobionts that are often elevated in Crohn’s disease.^15,16,18,48^

### Endocannabinoid-dependent proliferation of gut *E. coli* is mediated by the host cannabinoid receptor CB1

Endocannabinoids can directly impact bacterial populations by functioning as interkingdom hormones and modulating bacterial growth, physiology, and virulence potential.^42,46,49^ We previously reported that 2-AG antagonizes the bacterial histidine sensor kinase QseC, resulting in attenuated virulence and faster clearance of enteric pathogens from the gut.^46^ Because *qseC* is also encoded within the core *E. coli* genome ^50^, we next determined whether bacterial sensing of 2-AG contributes to the expansion of *E. coli* populations that occurs when endocannabinoid tone is increased. As observed with the parental strain, Magl inhibition stimulated significant proliferation of the *qseC*-deficient mutant (**Fig S2A**), thus ruling out this possibility. *In vitro* growth studies further showed that 2-AG does not directly impact *E. coli* growth kinetics (**Fig S2B-C**), suggesting that this endocannabinoid is unlikely being utilized as a nutrient to fuel *E. coli* proliferation. Together, these results led us to next interrogate the effects of endocannabinoid signaling in host tissues on intestinal *E. coli* populations.

Endocannabinoids exert their physiological effects on the host by engaging with cannabinoid receptors such as CB1 and CB2. To evaluate the relative contributions of each receptor, we tested whether co-treatment with a CB1 or CB2 antagonist counteracts the effects of Magl inhibition on gut *E. coli* expansion (**Fig 2A**). In groups co-treated with the CB1 antagonist, *E. coli* proliferation was no longer apparent in fecal or cecal contents (**Fig 2B-D**), suggesting that endocannabinoid stimulation of CB1 induces intestinal *E. coli* growth. To account for possible off-target effects of the CB1 antagonist, we repeated similar experiments in mice that lack the CB1 receptor (*Cnr1* KO) using heterozygous littermates as controls (*Cnr1* HET). Gut *E. coli* populations remained significantly unchanged following Magl inhibition in *Cnr1* KO mice (**Fig 2E-G**), thus confirming our initial results. Notably, acute CB1 antagonism or genetic CB1 deficiency did not impact basal fecal *E. coli* abundance (**Fig 2B**, **2E**). In contrast, significant *E. coli* expansion was still observed with Magl inhibition and CB2 antagonism (**Fig 2H-I**), suggesting that CB2 activity minimally contributes to this effect. Blocking CB1 or CB2 activity also prevented *Enterococcus* expansion following Magl inhibition (**Fig S2D-F**), highlighting a second reproducible effect on the microbiome that is dependent on host cannabinoid signaling. Altogether, our data strongly indicate that CB1 activation stimulates the proliferation of *E. coli* within the colon microbiome when host endocannabinoid activity is elevated.

**Fig. 2.**
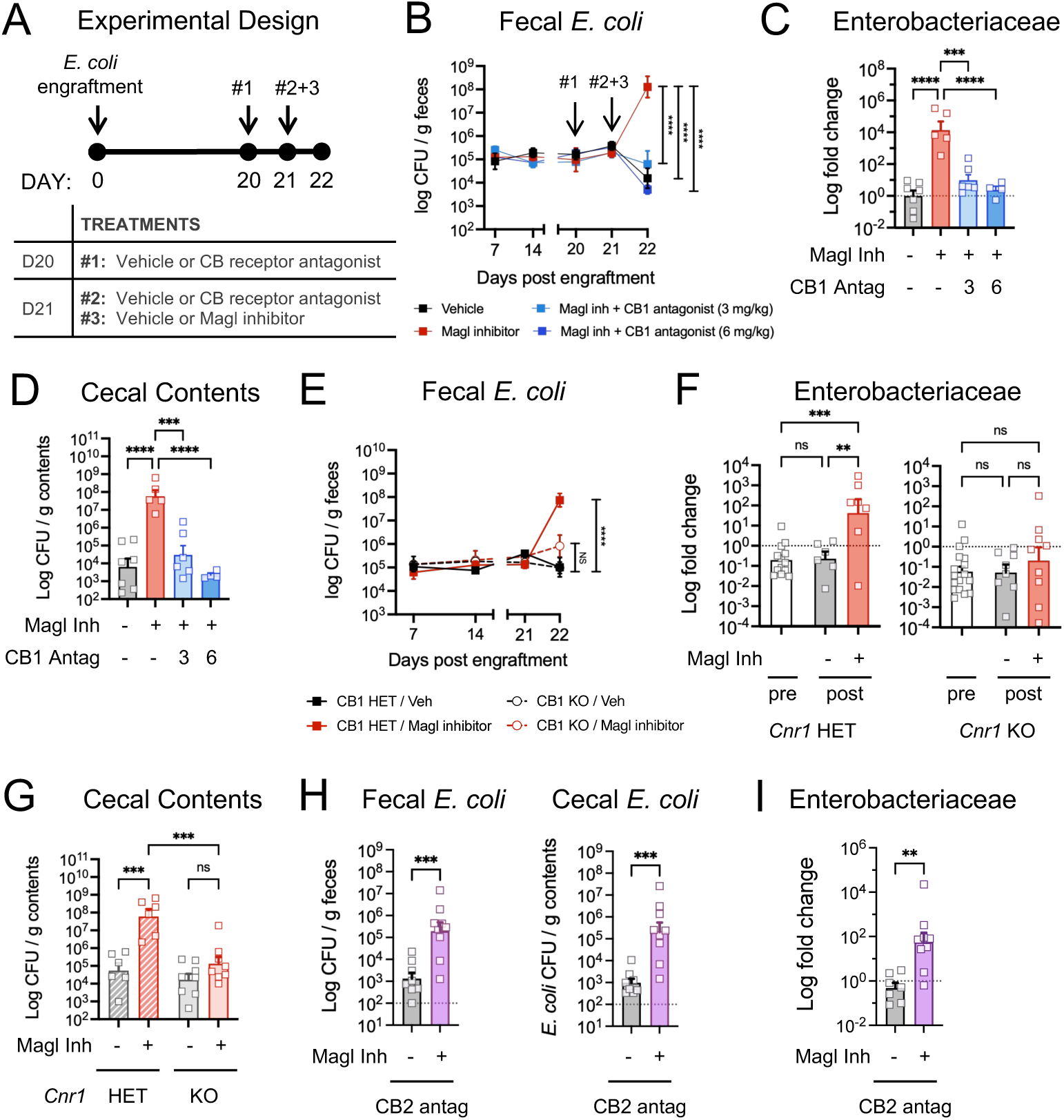
Endocannabinoid-dependent proliferation of gut *E. coli* is mediated by the host cannabinoid receptor CB1. A) Experimental design. B-C) Female mice engrafted with *E. coli* were treated with vehicle, Magl inhibitor, or CB1 antagonist in combination with Magl inhibitor (*n* = 4-7 per group). B) Fecal *E. coli* CFUs post engraftment. Two-way ANOVA with Sidak’s multiple comparison test. C) Post treatment relative abundance of fecal Enterobacteriaceae at day 22 relative to vehicle group as determined by targeted qPCR. One-way ANOVA with Tukey’s multiple comparisons test. D) Post treatment *E. coli* CFUs in cecal contents at day 22. One-way ANOVA with Tukey’s multiple comparisons test. E-G) Male and female *Cnr1* HET or KO mice were engrafted with *E. coli,* then treated with either vehicle or Magl inhibitor (*n* = 6-8 per group). E) Fecal *E. coli* CFUs post engraftment. Two-way ANOVA with Sidak’s multiple comparison test. F) Pre- and post-treatment relative abundance of fecal Enterobacteriaceae as determined by targeted qPCR. Data are represented as log fold change relative to pretreatment. One-way ANOVA with Tukey’s multiple comparison test. G) Post treatment *E. coli* CFUs in cecal contents. One-way ANOVA with Tukey’s multiple comparisons test. H-I) Male and female mice (*n* = 8-9 per group) were engrafted with *E. coli*, then treated with CB2 antagonist (5 mg/kg) alone or in combination with Magl inhibitor. H) Post treatment *E. coli* CFUs in feces and cecal contents. Student’s t test. I) Post treatment relative abundance of fecal Enterobacteriaceae relative to vehicle group as determined by targeted qPCR. Student’s t test. * *p* < 0.05, ** *p* < 0.01, *** *p* < 0.001, **** *p* < 0.000

### CB1 activity promotes signatures of microbial dysbiosis in a colitis-susceptible host

Having established that host endocannabinoid activity can reproducibly induce *E. coli* proliferation within the uninflamed gut, we next investigated the effects of CB1 signaling on microbiota composition in a host that is prone to developing chronic intestinal inflammation. *Il10-*deficient mice lack an immunomodulatory cytokine that is critical for maintaining mucosal homeostasis and are therefore highly susceptible to inflammation when exposed to specific microbial or environmental triggers.^51–53^ As observed in WT mice, Magl inhibition stimulated fecal and cecal *E. coli* proliferation in a CB1 dependent manner in *Il10* KO mice (**Fig 3A, S3A-B**). Moreover, 16S rRNA sequencing revealed significant CB1-dependent changes to the fecal microbiome with Magl inhibition in *Il10* KO mice (PERMANOVA: vehicle vs Magl inhibitor: *F* = 3.83, *p* = 0.001; Magl inhibitor vs combination treatment: *F* = 4.31, *p* = 0.002) while not impacting alpha diversity (**Fig 3B, S3C**). Similar to WT mice, Magl inhibition increased the relative abundance of Enterobacteriaceae, Christensenellaceae, and Anaerovoracaceae in a CB1-dependent manner (**Fig 3C, Table S1**). However, in contrast to WT mice, Ruminococcaceae and Enterococcaceae remained unchanged. Instead, Magl inhibition promoted the expansion of Staphylococcaceae, another low abundant family of facultative anaerobes. Magl inhibition also significantly impacted the three most abundant families in the *Il10* KO microbiome. This included the CB1-dependent expansion of Lactobacillaceae and Muribaculaceae and the CB1-dependent contraction of Lachnospiraceae, which includes many butyrate producers that are decreased in Crohn’s disease.^8,16,18^ Targeted qPCR analyses confirmed that Lachnospiraceae contraction reproducibly occurs specifically in *Il10* KO mice following Magl inhibition, and the CB1 dependent nature of this microbiome alteration (**Fig S3D-E**). Thus, in addition to its consistent effects on gut *E. coli* populations, our results suggest that acute CB1 activity in an inflammation susceptible host confers more extensive changes to microbial community structure that are frequently associated with inflammation-associated microbial dysbiosis.

**Fig 3.**
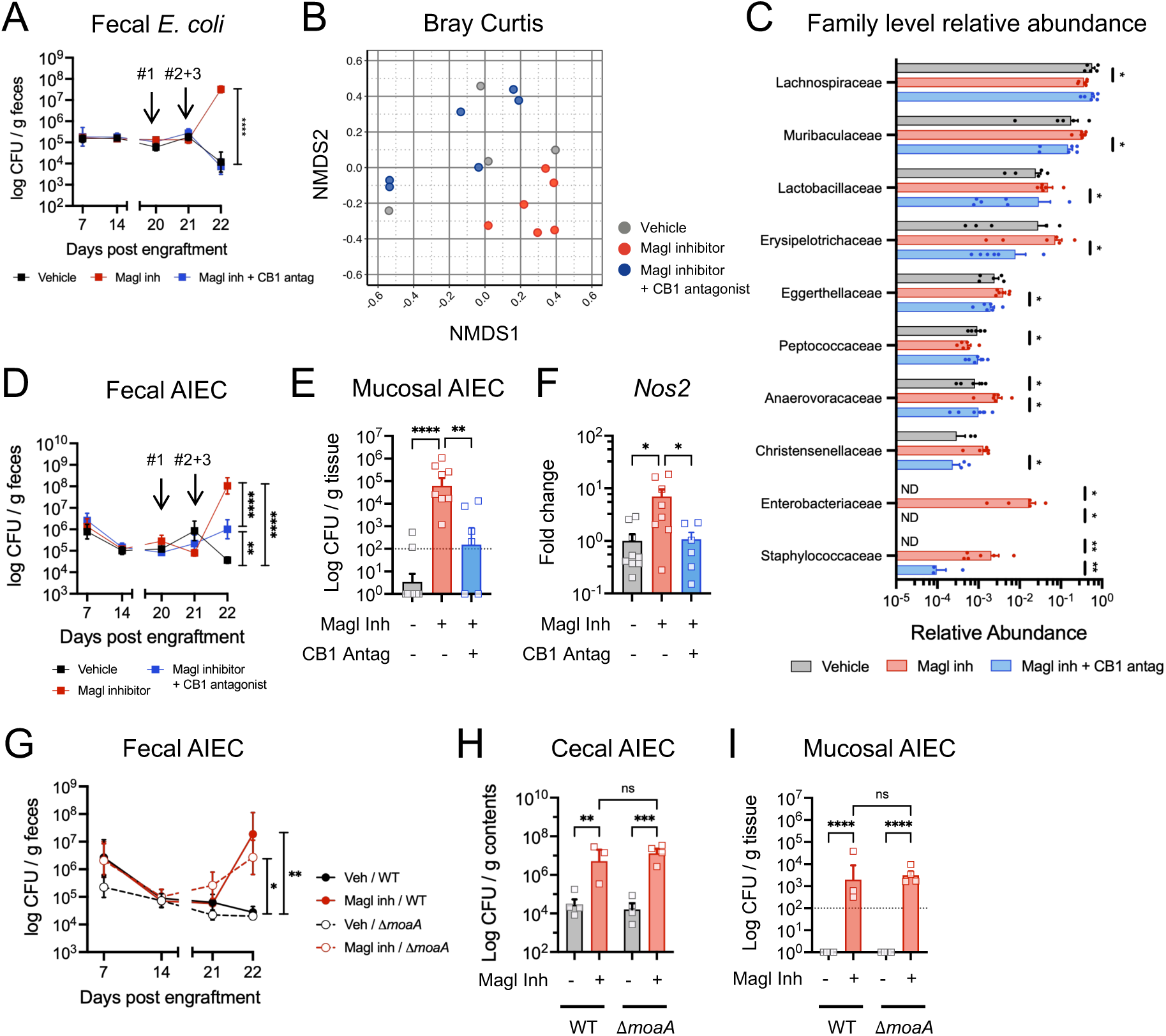
CB1 activity induces features of inflammation-associated dysbiosis in a colitis-susceptible host. A-C) Female *Il10* KO mice (n = 5-6 per group) were engrafted with *E. coli,* then treated with Magl inhibitor (32 mg/kg), or CB1 antagonist (3 mg/kg) in combination with Magl inhibitor following the experimental design in panel 2A. A) Fecal *E. coli* CFUs post engraftment. Two-way ANOVA with Sidak’s multiple comparison test. B) NMDS analysis of fecal microbiomes post-treatment. C) Relative abundance of bacteria families that were significantly altered with Magl inhibition relative to vehicle and/or the combination treatment. Datapoints were excluded if below detection. ND, not detected in any individual in the treatment group. D-F) Female *Il10* KO mice (n = 6-9 per group) were engrafted with AIEC, then three weeks later were treated with vehicle, Magl inhibitor (32 mg/kg), or CB1 antagonist (6 mg/kg) in combination with Magl inhibitor following experimental design in panel 2A. D) Longitudinal fecal AIEC CFUs. Two-way ANOVA with Sidak’s multiple comparisons test. E) Post treatment AIEC CFUs in colon mucus at day 22. F) Host expression of *Nos2* in colon tissues relative to vehicle. One-way ANOVA with Tukey’s multiple comparisons test. G-I) Female *Il10* KO mice (n = 3-4 per group) were engrafted with AIEC WT or Δ*moaA,* then three weeks later were treated with a single dose of vehicle or Magl inhibitor (32 mg/kg). G) Longitudinal fecal AIEC culture. Two-way ANOVA with Sidak’s multiple comparison test. H-I) Post treatment AIEC CFUs in H) cecal contents and I) colon mucus at day 22. One-way ANOVA with Tukey’s multiple comparisons test. * *p* < 0.05, ** *p* < 0.01, *** *p* < 0.001, **** *p* < 0.0001.

Adherent invasive *Escherichia coli* (AIEC) are opportunistic pathogens frequently associated with Crohn’s disease.^54^ AIEC can be innocuously present at low levels within the lumen of uninflamed hosts. However, their overgrowth in the inflamed gut facilitates mucosal niche formation, epithelial invasion, and translocation into the lamina propria, which can contribute to disease pathogenesis in inflammation susceptible hosts including *Il10* KO mice.^19^ We therefore investigated whether acute CB1 activity can promote AIEC infection of the mucosa in *Il10* KO mice. Consistent with our previous results, Magl inhibition induced luminal AIEC proliferation in a CB1-dependent manner, which corresponded with AIEC mucosal infection and increased *Nos2* expression in colon tissues (**Fig 3D-F, S3F**). *Lcn2* and *Tnfa* were also upregulated following Magl inhibition, albeit in a CB1-independent manner (**Fig S3G**). Collectively, these findings demonstrate that endocannabinoid signaling compromises colonization resistance normally exerted on resident AIEC, thus supporting their mucosal localization and elevating gut inflammatory markers in inflammation-susceptible hosts.

In the inflamed gut, the luminal availability of nitrate can increase as a result of host Nos2 activity, which can stimulate the overgrowth of *E. coli* via bacterial nitrate respiration.^17^ Considering that the CB1-dependent proliferation of AIEC coincides with increased *Nos2* expression, we next sought to investigate whether this effect is driven by bacterial nitrate respiration. The *E. coli* genome encodes three nitrate reductases, which require the cofactor molybdenum to function.^17^ We therefore first constructed a *moaA* deletion mutant that cannot synthesize molybdenum nor grow via nitrate respiration (**Fig S3H-I**).^17,55^ Genetic inactivation of MoaA-dependent metabolism failed to attenuate AIEC luminal proliferation and mucosal infection in *Il10* KO mice treated with the Magl inhibitor (**Fig 3G-I**). This suggests that nitrate respiration and other MoaA-dependent pathways unlikely contribute to *E. coli* proliferation when host endocannabinoid activity is acutely increased. Furthermore, the induction of *Nos2* is likely the result of endocannabinoid-dependent expansion of *E. coli* in an inflammation susceptible host.

### CB1 receptor activity induces signatures of metabolic stress in the colonic mucosa

We next investigated how host CB1 signaling alters the gut environment to support the rapid proliferation of gut *E. coli*. To accomplish this, we performed bulk RNAseq on colon mucosal tissues collected at three hours post Magl inhibition in the presence or absence of the CB1 antagonist. This time point was chosen because colon 2-AG levels are elevated with Magl inhibition (**Fig S4A**), but precedes detectable fecal *E. coli* proliferation (**Fig 1B**). We used a pharmacological approach to avoid possible basal changes to the host transcriptome that may occur due to lifelong *Mgll* or *Cnr1* deficiency. Furthermore, we utilized mice that lack Enterobacteriaceae in their microbiome to exclude possible effects of early *E. coli* expansion on host mucosal responses.

Gene set enrichment analysis (GSEA) revealed numerous pathways that were significantly enriched with Magl inhibition (**Fig 4A, S4B, Table S2**). Within the Hallmarks database, six shared pathways were positively enriched, and one shared pathway was negatively enriched, following Magl inhibition compared to the vehicle and combination treatment groups (**Fig 4A**). The top three positively enriched pathways in both comparisons included interferon signaling and oxidative phosphorylation (OXPHOS). Within the KEGG database, seven shared pathways were positively enriched, and one shared pathway was negatively enriched, following Magl inhibition compared to the other treatment groups (**Fig S4B**). OXPHOS represented the top positively enriched pathway with Magl inhibition in both comparisons. Notably, retrograde endocannabinoid signaling was also positively enriched with Magl inhibition, which is consistent with elevated endocannabinoid activity that is expected when tissue concentrations of 2-AG are increased. In contrast, OXPHOS was negatively enriched following combined Magl inhibition and CB1 antagonism relative to the vehicle group (**Fig 4A, S4B**). This is consistent with inverse agonism reported for AM281 in addition to its established antagonist activity at the CB1 receptor.^56^

**Fig 4.**
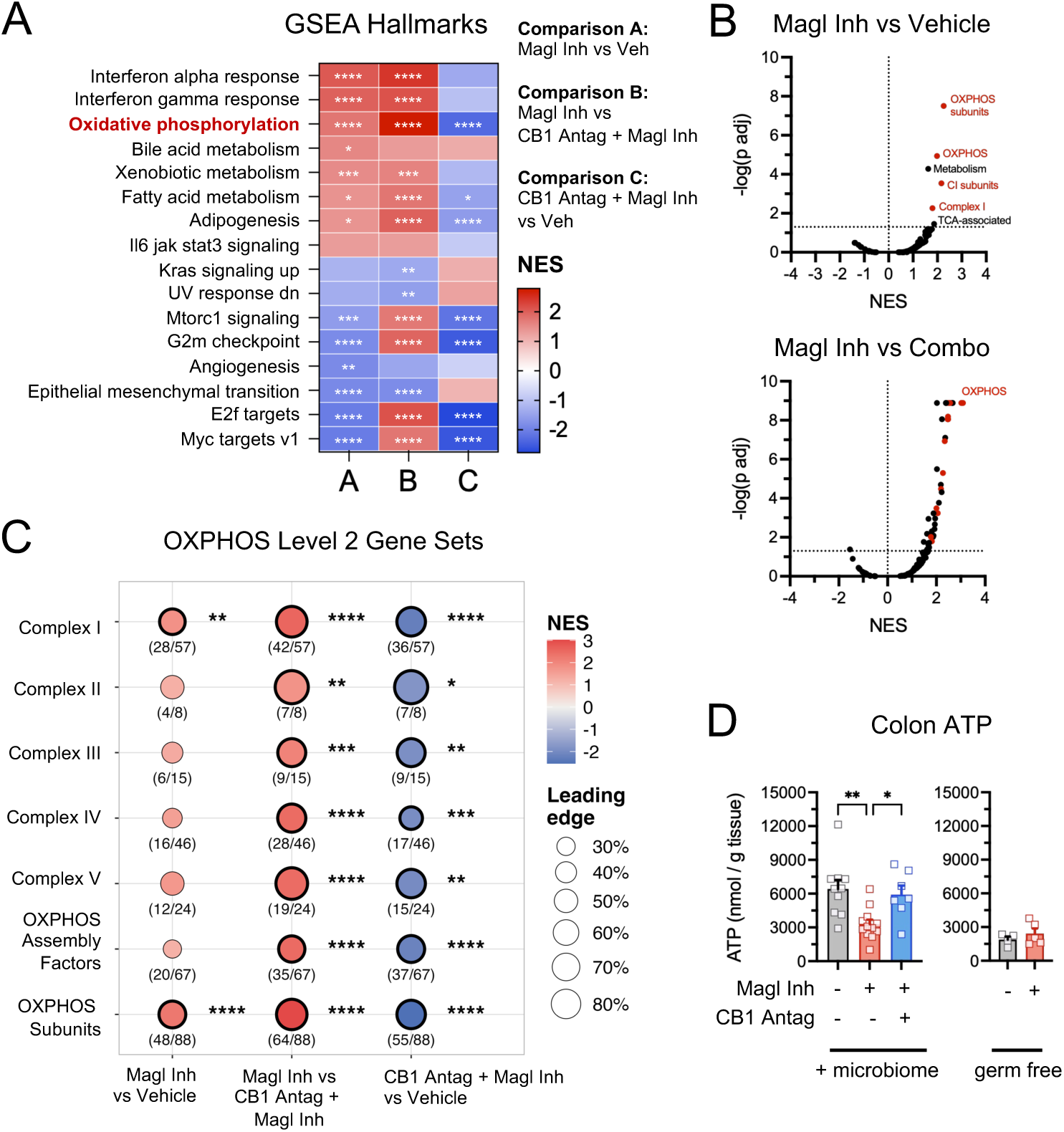
CB1 activity induces signatures of metabolic stress in the colonic mucosa. A-C) Female WT mice (n = 6-7 per group) were treated with a single dose of vehicle, Magl inhibitor (32 mg/kg), or CB1 antagonist (6 mg/kg) in combination with Magl inhibitor. Colon mucosal tissues were collected 3 hours post-treatment for bulk RNA-seq. A) Gene set enrichment analysis (GSEA) using the Hallmarks gene set collection. Only enriched pathways with FDR < 0.2 in comparisons A and B were included. * FDR < 0.05, ** FDR < 0.01, *** FDR < 0.001, **** FDR < 0.0001. B) Volcano plots of GSEA using the Mouse MitoCarta 3.0 Pathways collection. *P-*values are adjusted by Benjamini-Hochberg method (FDR). MitoPathways Levels 1, 2, and 3 are shown regardless of nesting. Red dots denote gene sets nested within Level 1 OXPHOS. Dashed lines represent NES = 0 and FDR = 0.05. C) Dot plot of GSEA results for Level 2 MitoPathways within the OXPHOS branch. Fill color represents the NES, while dot size corresponds with the leading-edge fraction, which is the proportion of detected genes in the gene set that fall within the leading edge. Numbers beneath each dot indicate the number of leading-edge genes over total genes detected in the gene set. * FDR < 0.05, ** FDR < 0.01, *** FDR < 0.001, **** FDR < 0.0001. D) Left: Male and female WT mice (*n* = 10-12 per group) engrafted with *E. coli* were treated with a single dose of vehicle, Magl inhibitor (32 mg/kg), or CB1 antagonist (6 mg/kg), in combination with the Magl inhibitor. Right: Female germ-free WT mice (*n* = 4-5 per group) were treated with a single dose of vehicle or Magl inhibitor (32 mg/kg). Data depict ATP concentrations in colon mucosal tissues at 12 hours post treatment. One-way ANOVA with Tukey’s post-test. * *p* < 0.05, ** *p* < 0.01

To further define the specific respiratory processes that are transcriptionally impacted by CB1 activity, we performed GSEA using the MitoCarta 3.0 database, a curated ontology of mitochondrial pathways consisting of proteins that localize to the mitochondria based on experimental evidence.^57^ OXPHOS was among the most positively enriched level 1 gene set following Magl inhibition relative to the other treatment groups (**Fig 4B, Table S2**). Moreover, among the six positively enriched MitoCarta gene sets in the Magl inhibitor group compared to vehicle, four are in the OXPHOS gene set. Among the 46 positively enriched MitoCarta gene sets in the Magl inhibitor group compared to the combination treatment group, 14 are in the OXPHOS gene set. The OXPHOS gene set includes genes that encode for assembly factors and structural components of the electron transport chain. Within OXPHOS, the level 2 gene sets OXPHOS Subunits and Complex I were positively enriched with Magl inhibition relative to the vehicle group, while gene sets associated with the other respiratory complexes were not significantly altered (**Fig 4B-C**). In contrast, level 2 gene sets associated with all complexes including OXPHOS assembly factors were positively enriched with Magl inhibition relative to the combination treatment. This corresponded with a significant negative enrichment of all level 2 gene sets within OXPHOS when comparing the combination treatment versus the vehicle, which is consistent with our GSEA results across the other databases. Notably, none of the leading genes within the OXPHOS gene sets in any comparison were mitochondrial encoded (**Table S2**). Together, our results demonstrate that acute CB1 activity induces the expression of nuclear encoded complex subunits and assembly factors of the electron transport chain.

The transcriptional upregulation of OXPHOS genes can be indicative of unmet cellular energy demands and mitochondrial stress due to diminished electron flow through the electron transport chain.^58,59^ Colonocytes derive the majority of their ATP from mitochondrial beta-oxidation and subsequent OXPHOS; therefore, ongoing mitochondrial dysfunction can decrease epithelial ATP production.^5^ Using ATP as a metabolic indicator of respiration, we found that Magl inhibition significantly decreased colon ATP levels in mucosal tissues in a CB1-dependent manner at 12 hours post treatment (**Fig 4D**), which coincides with peak fecal *E. coli* expansion (**Fig 1B**). In contrast, Magl inhibition had no effect on colon ATP levels in germ-free mice, which likely reflects the energy-deprived state of colonocytes in the absence of microbial-derived butyrate.^4,5^ Consistent with previous studies, colonic ATP production was generally lower in germ-free mice compared to mice colonized with a microbiota (**Fig 4D**).^60^ Taken together, our results indicate that endocannabinoid activity at the CB1 receptor decreases ATP production within the colonic mucosa, which may be indicative of diminished respiratory metabolism and disrupted mucosal bioenergetics.

### CB1 activity stimulates the aerobic expansion of gut *E. coli*

Respiration within the colon epithelium is an oxygen consuming process that is critical for maintaining the anoxic environment of the lumen.^4^ Disruption of OXPHOS is linked to microbial dysbiosis, including facilitating the overgrowth of facultative anaerobes via bacterial aerobic respiration.^4^ Our transcriptomics analyses and the decrease in mucosal ATP led us to hypothesize that stimulation of CB1 induces Enterobacteriaceae proliferation by facilitating the infusion of oxygen into the lumen. The *E. coli* genome encodes three terminal oxidases – cytochrome bo_3_ oxidase (CyoABCD), cytochrome bd-I oxidase (CydAB), and cytochrome bd-II oxidase (AppBC).^61–63^ Each are maximally expressed at different oxygenation levels and redox environments.^64–67^ To test whether aerobic respiration in *E. coli* is required for supporting its proliferation following Magl inhibition, we first investigated whether genes encoding each terminal oxidase are induced. To accomplish this, and ensure detection of *E. coli*-specific transcripts, we first mono-colonized germ-free mice with *E. coli* and then treated mice with either the Magl inhibitor or vehicle control (**Fig 5A**). Targeted expression analyses revealed that *cydA* – and not genes that encode subunits of the other terminal oxidases – was significantly induced at 12 hours following Magl inhibition (**Fig 5B**). Notably, the CydAB terminal oxidase is an established mediator of aerobic growth in *E. coli* when physiological hypoxia in the colon is disrupted.^4,68,69^ Remarkably, this coincided with a significant increase in luminal *E. coli* despite initial high population densities prior to treatment (**Fig 5C-D**), further reinforcing the idea that endocannabinoid signaling in host tissues induces *E. coli* proliferation.

**Fig 5.**
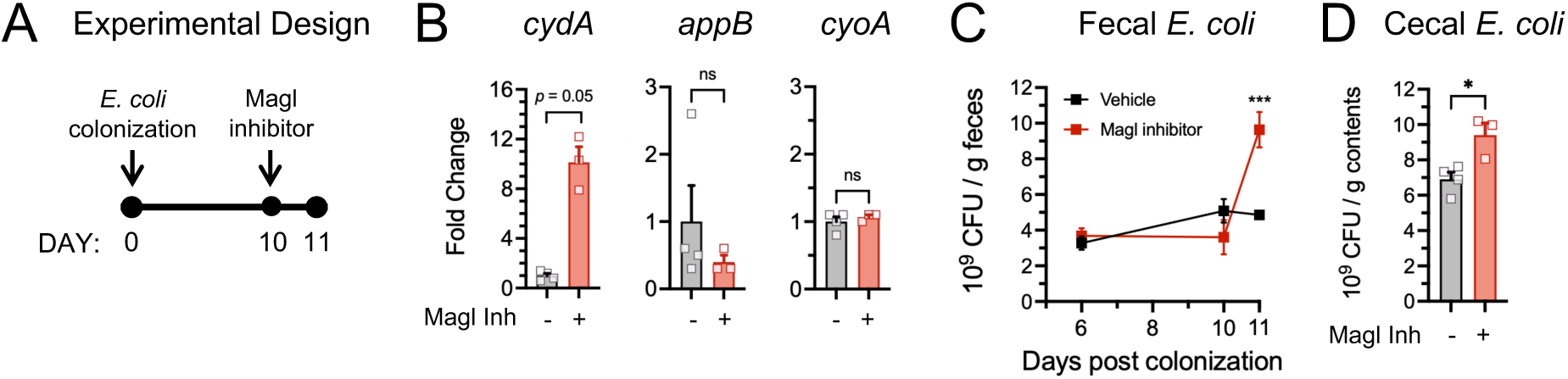
Expression of *cydA* in luminal *E. coli* is induced following Magl inhibition. Female germ-free mice (*n* -3-4 per group) were colonized with *E. coli*. Ten days later, mice were treated with a single dose of vehicle or Magl inhibitor (32 mg/kg). A) Experimental design. B) Relative expression of select *E. coli* genes in cecal contents. Mann-Whitney test. C) Fecal *E. coli* CFUs post colonization. Two-way ANOVA with Sidak’s multiple comparison test. D) Post-treatment *E. coli* CFUs in cecal contents. Student’s t test.

Guided by our expression data, we next tested whether deletion of *cydAB* abrogates *E. coli* expansion induced by Magl inhibition. To accomplish this, we performed *in vivo* growth competitions between the parental strain and *cydAB* deletion mutant utilizing the experimental design depicted in **Fig 6A**. We first confirmed that the antibiotic selection markers utilized to distinguish strains via selective culture do not impact *E. coli* colonization fitness following our treatments (**Fig 6B**). Using this approach, we found that Δ*cydAB* exhibited a competitive disadvantage relative to its parental strain when endocannabinoid levels were elevated (**Fig 6C, S5A**). Nonetheless, this mutant still proliferated following Magl inhibition -albeit to a lesser extent. Similarly, in our engraftment model (**Fig 1A**), proliferation of Δ*cydAB* was attenuated in mice treated with the Magl inhibitor (**S5C-D**). Together, these results indicate the involvement of other terminal oxidases or metabolic pathways that mediate *E. coli* expansion.

**Fig 6.**
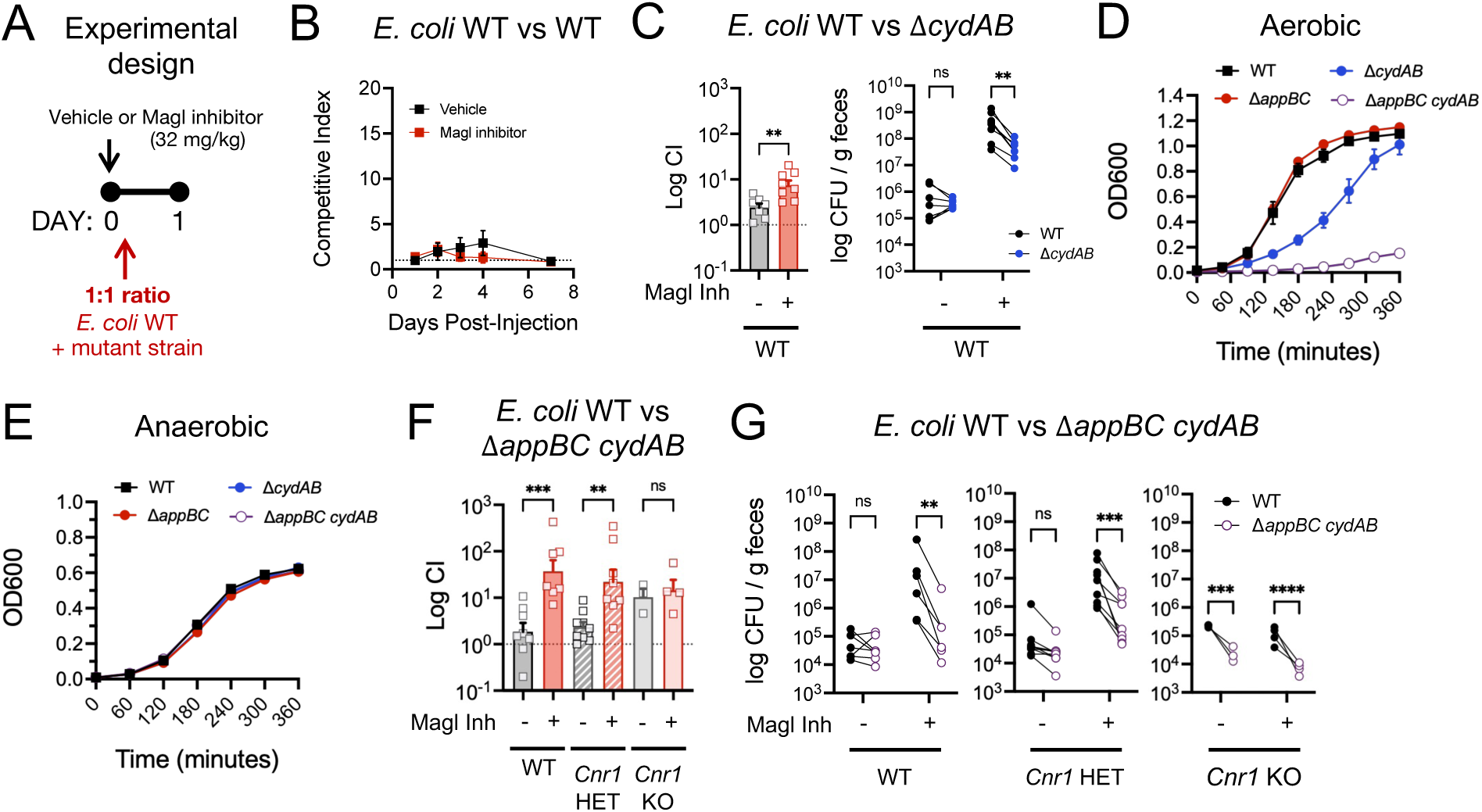
CB1 activity stimulates the aerobic expansion of gut *E. coli*. A) Experimental design of *in vivo* growth competitions. Mice were treated with vehicle or Magl inhibitor (32 mg/kg), then gavaged 2 hours later at a 1:1 ratio with the competing *E. coli* strains. B) Female mice (*n* = 4 per group) were gavaged with the parental *E. coli* strains, each tagged with a different antibiotic selection marker. Competitive indices (marker 1 vs marker 2) from fecal culture. C) Female WT mice (n = 7-8 per group) were gavaged with the indicated *E. coli* isogenic mutant and parental strain. Competitive indices (parental vs mutant) were calculated from fecal culture. Student’s t test (first panel); two-way ANOVA with Sidak’s multiple comparison test (second panel). D) Aerobic and E) anaerobic growth curves of individual *E. coli* strains. Data are combined from at least two independent experiments with 3 technical replicates per experiment. F-G) Male and female WT, *Cnr1* HET, or *Cnr1* KO mice (n = 3-8 per group) were gavaged with the indicated *E. coli* isogenic mutant and parental strain. F) Competitive indices (parental vs mutant) were calculated from G) *E. coli* CFUs cultured from feces. Two-way ANOVA with Sidak’s multiple comparison test. ** *p* < 0.01, *** *p* < 0.001, **** *p* < 0.0001

To delineate the possible contribution of the other terminal oxidases, we constructed a collection of single and double terminal oxidase mutants to empirically identify an *E. coli* mutant that exhibits the greatest growth defect under aerobic growth (**Fig 6D, S5B**). We identified one mutant (Δ*appBC cydAB*) that exhibited minimal growth in the presence of oxygen, but no growth defect anaerobically (**Fig 6D-E**). We next performed *in vivo* growth competitions between the parental and Δ*appBC cydAB* strains in mice treated with the vehicle or Magl inhibitor. In WT and *Cnr1* HET mice, the parental strain significantly outcompeted Δ*appBC cydAB* following Magl inhibition, while exhibiting no competitive disadvantage in vehicle treated mice (**Fig 6F-G, S5E-F**). Moreover, minimal expansion of Δ*appBC cydAB* was observed following Magl inhibition. In contrast, treatment specific differences in colonization fitness between *E. coli* strains were not observed in *Cnr1* KO mice. Interestingly, while no strain significantly expanded with Magl inhibition in *Cnr1* KO mice, in agreement with our earlier results (**Fig 2E-F**), the Δ*appBC cydAB* mutant consistently exhibited a fitness disadvantage relative to the parental strain regardless of treatment. This suggests that life-long *Cnr1* deficiency may impact basal luminal oxygen tensions within the colon. In our engraftment model, in contrast to the Δ*cydAB,* deletion of *appBC* did not attenuate *E. coli* proliferation in mice treated with the Magl inhibitor (**Fig S5C-D**), suggesting that this terminal oxidase may serve a compensatory role in the absence of functional CydAB. Finally, we also attempted to engraft WT mice with Δ*appBC cydAB;* however stable colonization of this mutant was not achieved in most mice (data not shown). Taken together, these data support our hypothesis that CB1-dependent *E. coli* expansion is primarily mediated by aerobic respiration.

### CB1 receptor activity exacerbates gut infection by an Enterobacteriaceae pathogen

Enteric pathogens deploy virulence factors to overcome colonization resistance and to establish replicative niches within host tissues.^2^ Enterobacteriaceae pathogens such as *Citrobacter rodentium* utilize their virulence factors to remodel the colonic epithelium, which facilitates their oxygen-dependent proliferation within the gut.^70^ Because *C. rodentium* relies on the CydAB terminal oxidase to maximally colonize the gut during early infection ^70^, we next investigated whether endocannabinoid activity can enhance *C. rodentium* infection (**Fig 7A**). As observed with *E. coli,* Magl inhibition enhanced *C. rodentium* infection in a dose dependent manner with near peak fecal shedding evident by day 2 versus day 6 post infection (**Fig 7B, S6A**). Pathogen infection of cecal tissues at day 6 was significantly elevated with Magl inhibition, which corresponded with increased *Nos2* expression and exacerbated cecal pathology characterized by marked hyperplasia and severe edema (**Fig 7C-H**). Notably, cecal *Nos2* expression strongly correlates with tissue pathogen burdens and the extent of pathology in this model.^46^ In contrast, repeated Magl inhibition did not induce cecal pathology or *Nos2* expression in uninfected mice that also lack resident Enterobacteriaceae (**Fig 7D, H**).

**Fig. 7.**
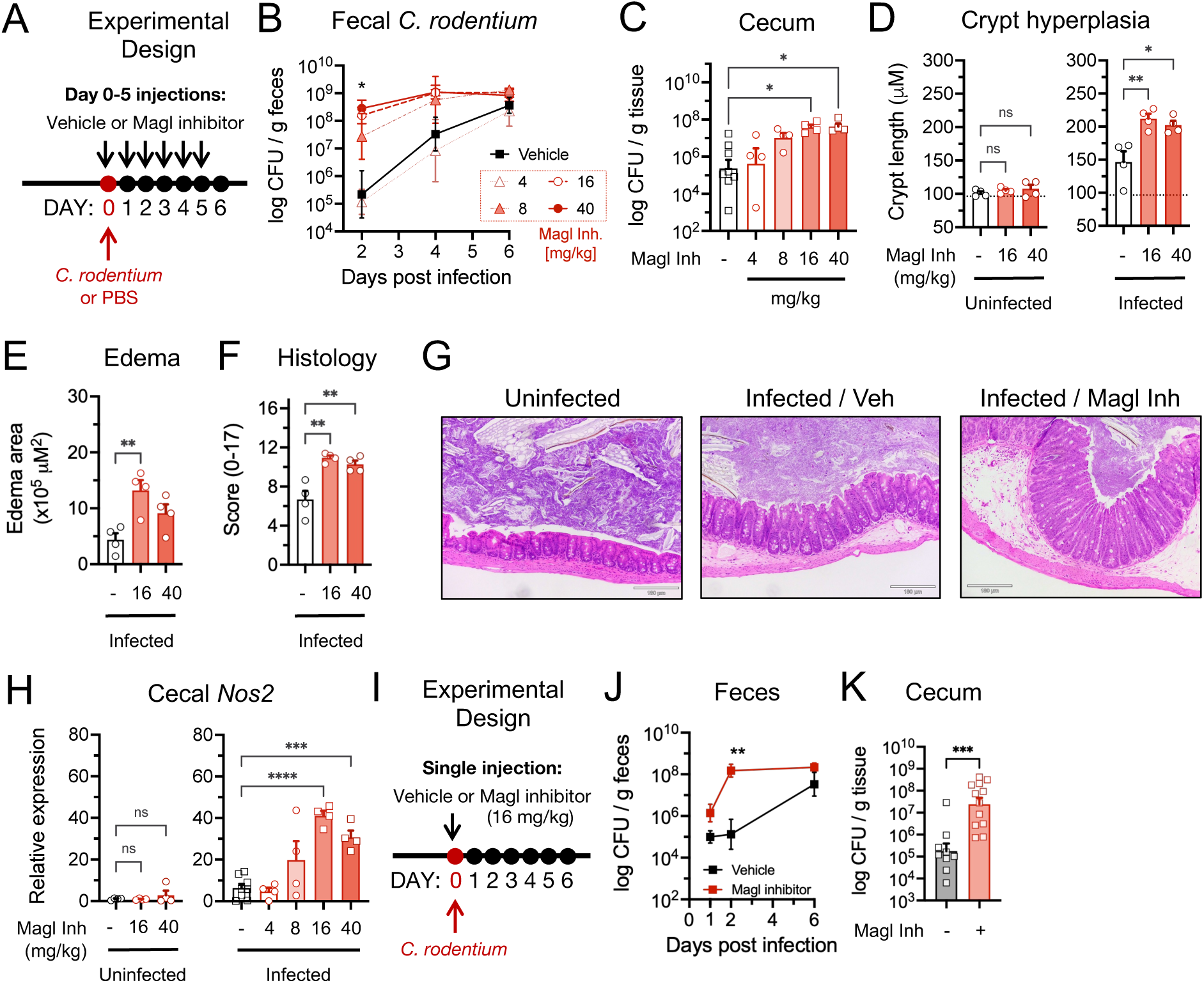
Host endocannabinoid activity during early infection exacerbates colitis. A-G) Female mice were infected with *C. rodentium* and administered daily injections of vehicle or Magl inhibitor. A) Experimental design. B) Fecal *C. rodentium* CFUs (*n* = 4-8 per group, pooled from 2 cohorts). Two-way ANOVA with Sidak’s multiple comparisons test. * *p* < 0.05 in vehicle versus Magl inhibitor at 16 and 40 mg/kg. C) *C. rodentium* CFUs in cecal tissues at day 6. One-way ANOVA with Tukey’s multiple comparisons test. D) Crypt hyperplasia length measurements, E) edema lesion area, and F) histology scores of cecal sections at day 6 (*n* = 4 per group from 1 cohort). Dotted line represents average crypt length in uninfected, vehicle-treated mice. One-way ANOVA with Tukey’s multiple comparisons test. G) Representative histology of ceca at day 6. Dose of Magl inhibitor was 16 mg/kg. H) Relative expression of *Nos2* in cecal tissue at day 6 relative to vehicle treated, uninfected controls (*n* = 4-8 per group, pooled from 2 cohorts). One-way ANOVA with Tukey’s multiple comparisons test. I-K) Female mice (*n* = 9-12 per group, pooled from 2 cohorts) were administered a single dose of vehicle or Magl inhibitor, then two hours later were infected with *C. rodentium.* Two mice in the vehicle group were excluded because they remained uninfected throughout duration of experiment. I) Experimental design. J) Fecal *C. rodentium* CFUs. Two-way ANOVA with Sidak’s multiple comparison test. K) *C. rodentium* CFUs in cecal tissues at day 6. Student t-test. * *p* < 0.05, ** *p* < 0.01, *** *p* < 0.001, **** *p* < 0.0001

Considering the accelerated *C. rodentium* infection dynamics observed with repeated Magl inhibition, we next tested whether acute Magl inhibition prior to pathogen challenge is sufficient to induce rapid early *C. rodentium* colonization and impact later infection of host tissues (**Fig 7I**). Acute Magl inhibition recapitulated the accelerated infection dynamics of *C. rodentium* observed with repeated treatments (**Fig 7J**), which resulted in elevated pathogen burdens in cecal tissues six days later (**Fig 7K**). Finally, considering our observations with gut *E. coli* (**Fig 1-2**), we tested whether increased endocannabinoid activity can rescue the colonization defect of an attenuated *C. rodentium* mutant. Using a *fadL* deficient strain, which we previously reported as having a low infection rate ^46^, we found that repeated treatment with the Magl inhibitor rescued its colonization defect (**Fig S6B**). Notably, FadL is also required for 2-AG sensing ^46^, thus suggesting that the effects of Magl inhibition on *C. rodentium* colonization occur independently from the anti-virulence effects of 2-AG that become apparent during the peak and resolution stages of infection.^46^

We next tested whether the effects of host endocannabinoid activity on *C. rodentium* infection are also CB1 dependent (**Fig 8A**). We found that CB1 antagonism prevented the early increase in *C. rodentium* fecal shedding observed at day 2 with Magl inhibition (**Fig 8B**). This corresponded with reduced pathogen burdens in cecal tissues, decreased *Nos2* induction, and attenuated tissue damage by day 6 post infection (**Fig 8C-G**). Considering the elevated fecal abundance of *C. rodentium* at day 2, we next investigated whether CB1 activity also enhances pathogen infection of cecal tissues at this time point. In mice treated acutely with the Magl inhibitor, CB1 antagonism significantly reduced *C. rodentium* colonization of cecal tissues at day 2 (**Fig 8H**), suggesting that earlier luminal expansion permits faster mucosal infection. Notably, despite being present in fecal samples, *C. rodentium* was not detectable in cecal tissues with CB1 antagonism alone, suggesting that basal CB1 activity may facilitate early pathogen colonization of cecal tissues. In support of this idea, relative endocannabinoid concentrations in cecal tissues were significantly increased during early infection compared to uninfected controls (**Fig 8I**). Finally, we tested whether acute CB1 activation using a synthetic agonist can recapitulate the effects observed with Magl inhibition. Cannabinoid receptor agonism significantly increased *C. rodentium* colonization of cecal tissues at day 2 post infection at levels comparable to treatment with the Magl inhibitor (**Fig 8J**). Collectively, our results demonstrate that CB1 activity enables aerobically respiring pathogens to rapidly overcome colonization resistance in the non-inflamed gut, resulting in accelerated infection dynamics and exacerbated disease.

**Fig. 8.**
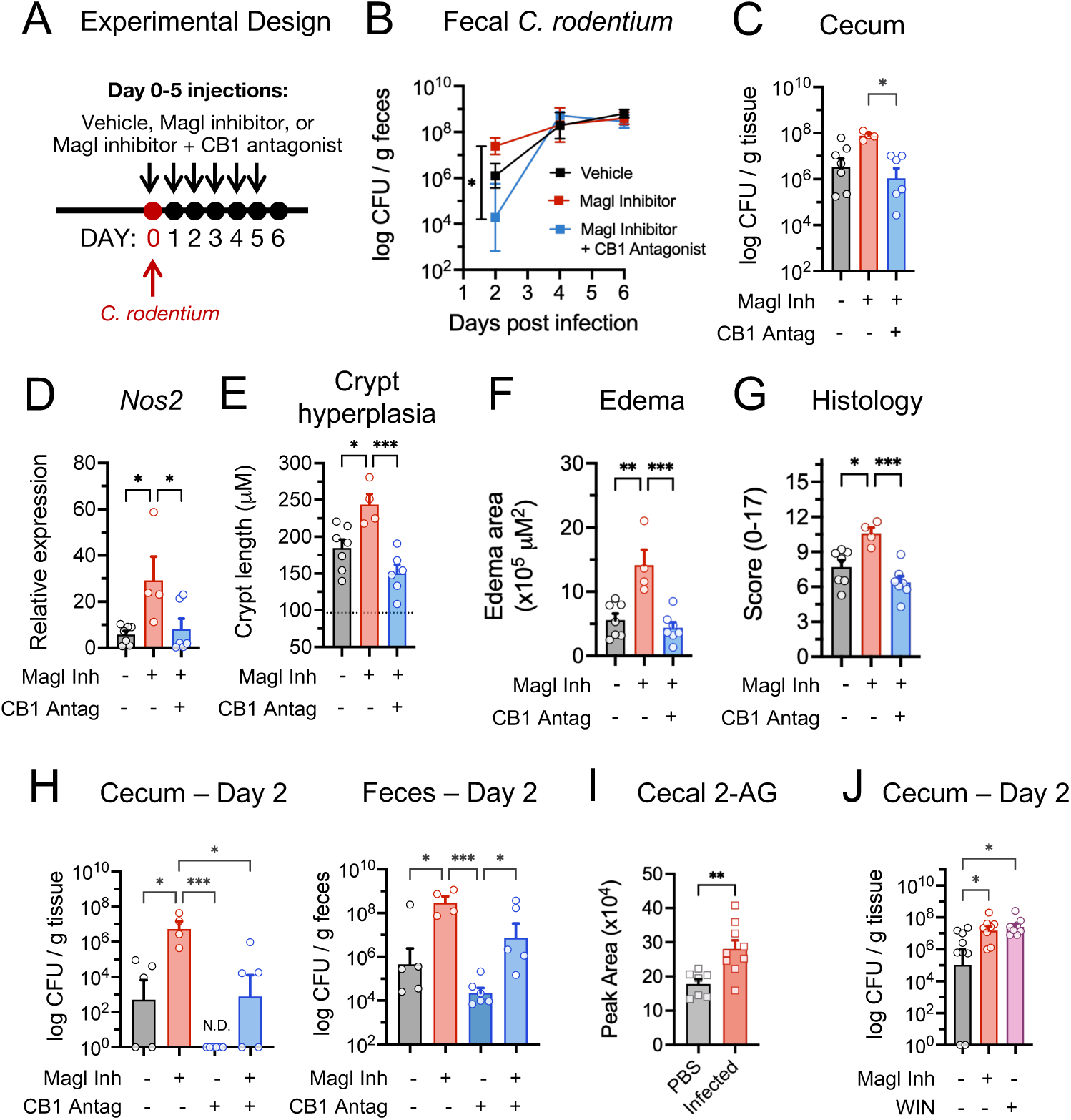
CB1 receptor activity enhances early infection by an Enterobacteriaceae pathogen. A-F) Female mice (*n* = 4-7 per group) were infected with *C. rodentium* and administered daily injections of vehicle, Magl inhibitor (16 mg/kg), or CB1 antagonist (3 mg/kg) in combination with Magl inhibitor. A) Experimental design. B) Longitudinal fecal *C. rodentium* CFUs. Two-way ANOVA with Sidak’s multiple comparisons test. C) *C. rodentium* CFUs in cecal tissues at day 6 post infection. One-way ANOVA with Tukey’s multiple comparisons test. D) Relative expression of *Nos2* in cecal tissue at day 6 relative to vehicle treated, uninfected controls. One-way ANOVA with Tukey’s multiple comparisons test. E) Crypt hyperplasia measurements, F) edema lesion area, and G) histology scores of cecal sections at day 6 post infection. Dotted line represents average crypt length in cecum of a non-infected, vehicle treated WT mouse. One-way ANOVA with Tukey’s multiple comparisons test. H) Female mice (*n* = 4-6 per group) were infected with *C. rodentium* and administered daily injections of vehicle, Magl inhibitor, (16 mg/kg), CB1 antagonist (3 mg/kg), or both. *C. rodentium* CFUs in cecal tissues and feces at day 2 post infection. One-way ANOVA with Tukey’s multiple comparisons test. ND, not detected. I) Female mice (*n* = 7-9 per group, pooled from 2 cohorts) were gavaged with PBS or *C. rodentium*. Detection and quantification of 2-AG by LC-MS in cecal tissues at day 3 post infection. Student’s t test. J) Female mice (*n* = 8-9 per group) were infected with *C. rodentium* and administered daily injections of vehicle, Magl inhibitor (16 mg/kg), or the CB1/CB2 agonist WIN55,212-2 (5 mg/kg). *C. rodentium* CFUs in cecal tissues at day 2 post infection. One-way ANOVA with Tukey’s multiple comparisons test. * *p* < 0.05, ** *p* < 0.01, *** *p* < 0.001.

## Discussion

Clinical studies have reported altered endocannabinoid tone in patients with inflammatory bowel diseases (IBD) including Crohn’s disease.^35–41^ This includes a positive correlation between elevated endocannabinoids and microbial dysbiosis in active Crohn’s disease, an association not observed in quiescent disease or other IBD sub-types.^42–44^ However, it remains unclear whether the endocannabinoid system can contribute to disease pathogenesis, which is critical for investigating its potential as a therapeutic target. Here, we demonstrate that stimulating host endocannabinoid activity at the CB1 receptor induces acute metabolic injury within the colon mucosa, resulting in the aerobic expansion of gut Enterobacteriaceae. Our data further suggest that CB1 activity disrupts mucosal respiration, thus diminishing host consumption of oxygen. This metabolic perturbation is consistent with Enterobacteriaceae overgrowth fueled by the diffusion of unconsumed oxygen into the lumen. Taken together, our findings establish host endocannabinoid activity as a mediator of microbial dysbiosis through its perturbation of mucosal metabolism, which can impact host risk for intestinal disease.

Animal models have been instrumental in identifying putative links between the host endocannabinoid system, mucosal immunity, and the gut microbiome.^20,21^ This includes changes in microbiome composition in hosts subjected to chronic alterations in endocannabinoid activity while consuming a high fat diet.^71,72^ Conversely, germ-free states and diet-induced changes to the microbiome are associated with altered host endocannabinoid tone.^73–75^ In rodent models of chemically induced colitis, *Cnr1* deficiency or CB1 blockade during inflammation development exacerbates disease, while increasing endocannabinoid levels is generally protective.^29–35^ Adding to this body of work, our findings demonstrate that acute stimulation of CB1 activity under homeostasis induces changes to the colon microbiome that recapitulate common features of inflammation associated dysbiosis. Most notably, this includes the overgrowth of pathogenic and commensal Enterobacteriaceae. The subsequent effects on the host are dependent on colitis susceptibility and Enterobacteriaceae virulence potential. This highlights the complexity of the enteric endocannabinoid system where the impact on inflammation outcomes clearly depends on the mechanism and duration of endocannabinoid manipulation and the stage of intestinal disease. Consistent with this idea, intestinal and systemic endocannabinoid tone differ among IBD subtypes and between quiescent versus active disease states.^35–41^

In the context of infectious colitis, acute CB1 stimulation in an inflammation resistant host is sufficient for an Enterobacteriaceae pathogen to rapidly overcome initial colonization resistance, thus accelerating mucosal infection and exacerbating disease. Aerobic respiration primarily supports *C. rodentium* proliferation during early infection ^70^, which is consistent with the idea that CB1-dependent oxygen infusion may accelerate these dynamics. However, we previously demonstrated that the endocannabinoid 2-AG also functions as an anti-virulence signal in *C. rodentium* and other enteric pathogens.^46^ Furthermore, in *Mgll*-deficient mice in which colon 2-AG is chronically elevated, challenge with *C. rodentium* results in decreased tissue damage and mucosal pathogen burdens at peak infection, followed by faster pathogen clearance.^46^ Interestingly, early infection dynamics are not impacted in these mice compared to *Mgll*-sufficient controls when the effects of acute Magl inhibition on early pathogen colonization are apparent. Notably, chronic Magl deficiency promotes CB1 desensitization as a result of prolonged receptor stimulation by 2-AG.^76^ This may explain the lack of differences in pathogen colonization during early infection when comparing *Mgll*-sufficient and *Mgll*-deficient mice, while possibly augmenting the anti-virulence effects of 2-AG when host CB1 activity is diminished. Indeed, our data show that CB1 antagonism alone is sufficient to decrease fecal *C. rodentium* burdens and inhibit mucosal colonization during early infection, which corresponds with elevated 2-AG concentrations. These results suggest that the onset of inflammation may naturally elevate local endocannabinoid concentrations, which in turn may facilitate early pathogen colonization by disrupting host respiratory metabolism via CB1 signaling. More broadly, our findings highlight the importance of host endocannabinoid tone as a potential predictor of inflammation outcomes in intestinal disease. Thus, it will be critical to establish how endocannabinoid levels and cannabinoid receptor expression, desensitization, and activity are modulated during inflammation development and resolution in specific disease contexts.

In Crohn’s disease, the development of disease is driven in part by an inability to appropriately respond to acute injury to the mucosa, which can lead to sustained tissue damage and chronic inflammation. In conjunction, microbial dysbiosis contributes to disease pathogenesis. Here, we show that in *Il10* KO mice that are genetically susceptible to developing chronic colitis, acute CB1 activity stimulates the proliferation and mucosal localization of an *E. coli* pathobiont. This corresponds with the significant induction of mucosal inflammatory markers that are commonly associated with active Crohn’s disease.^77^ More substantial changes to the microbiota were also observed in *Il10* KO mice, including the CB1-dependent contraction of oxygen-sensitive Lachnospiraceae, key producers of butyrate that fuels epithelial beta-oxidation and OXPHOS to help maintain mucosal homeostasis.^8^ Indeed, Lachnospiraceae are often decreased in active Crohn’s disease in conjunction with Enterobacteriaceae enrichment.^16,18^ While it is tempting to speculate that luminal oxygen influx may have contributed to the depletion of Lachnospiraceae in *Il10* KO mice, similar changes were curiously not observed in inflammation-resistant WT mice despite comparable aerobic expansion of Enterobacteriaceae. The mechanisms underlying this enhanced microbiome resiliency observed in WT versus *Il10* KO mice in the context of endocannabinoid signaling is worthy of exploring in future studies. Taken together, our data suggest that CB1 stimulation transiently disrupts mucosal homeostasis and alters the microbiome to a more proinflammatory configuration. This acute injury to both host and microbiome may then contribute to disease development or progression in a host that is susceptible to intestinal inflammation.

Mitochondrial dysfunction within intestinal tissues is a well-established feature of IBD.^1^ Moreover, in mouse models of Crohn’s disease, mitochondrial dysfunction often precedes intestinal epithelial damage and inflammatory disease.^78–80^ Notably, in extraintestinal tissues such as neurons and myocardial muscle, CB1 signaling has been linked to mitochondrial dysfunction including decreased OXPHOS.^81–84^ Our findings indicate that CB1 activity may also compromise mitochondrial function in the colonic mucosa, resulting in decreased ATP production and host oxygen consumption. Our transcriptomics analyses reveal that acute stimulation of CB1 activity promotes the upregulation of electron transport chain complexes, especially subunits associated with Complex I. Interestingly, acute treatment with established Complex I inhibitors also induces the expression of genes associated with the electron transport chain and OXPHOS.^59,85,86^ Such upregulation of respiratory genes is a putative signature of acute mitochondrial stress that may reflect the attempt of cells to compensate for impaired electron transport to address unmet energy demands.^58^ In agreement with this idea, our data show that mucosal ATP production is significantly diminished with CB1 activity. However, this response is distinct from the metabolic remodeling that occurs with chronic mitochondrial dysfunction in inflamed tissues including the sustained downregulation of OXPHOS reported in active IBD.^18,86–88^ Thus, it will be interesting to investigate in future studies whether prolonged CB1 activity also promotes chronic metabolic dysfunction especially in the context of intestinal inflammation.

In summary, our findings have identified the endocannabinoid system as a critical host signaling network that influences mucosal bioenergetics and luminal oxygenation within the gut. In the context of Crohn’s disease, endocannabinoid signaling may impair respiratory metabolism within mucosal tissues, disrupt physiological hypoxia within the colon epithelium, and promote microbial dysbiosis via luminal oxygen flux, which together can contribute to disease. However, future studies are needed to establish whether the increase in endocannabinoid tone reported in some Crohn’s disease patients translates to increased intestinal CB1 activity and downstream disruptions in mitochondrial function, epithelial metabolism, and microbial symbiosis. Nonetheless, our findings reveal a potential therapeutic entry point for modulating mucosal bioenergetics and luminal oxygenation to limit microbial dysbiosis and risk for intestinal inflammation especially in individuals susceptible to IBD.

## Resource Availability

16S rRNA sequencing data (accession number PRJNA1425032) and host RNAseq data (accession number PRJNA1430417) have been deposited at the Sequence Read Archive. All datasets will be publicly available as of the date of publication.

## Acknowledgements

This work was supported by the following funding sources: NIH/NIAID R01 AI84916-01 (ME); NIH/NIGMS COBRE Center for Dietary Supplements and Inflammation 5P20GM103641-01 (PIs: Drs. Mitzi and Prakash Nagarkatti; Target Faculty Grant awarded to ME); NIH/NIGMS SC INBRE P20GM103499 (SIRP grant awarded to MKM); USC SPARC Graduate Research Grant (MKM); USC Magellan Undergraduate Research Grants (MTA, CAK, AEB); USC Honors College Grant (MTA, CAK). We acknowledge the following Core facilities: Histopathology Core at UT Southwestern Medical Center for tissue processing and staining for histopathology analysis; University of Alabama Birmingham Microbiome Core for the 16S rRNA sequencing and initial bioinformatics analysis; qPCR was performed by the University of South Carolina Functional Genomics Core, supported by NIGMS P30GM154632 award (RRID:SCR_026178); LC-MS analyses were performed by the University of South Carolina Mass Spectrometry Facility, supported by NSF-MRI Program Award 1828059; and Duke University School of Medicine for the use of the Sequencing and Genomic Technologies Shared Resource, which provided the RNA sequencing services.

## Declaration of interests

The authors declare no competing interests.

## Supplemental Information

Document S1. Figures S1–S6, Supplementary Materials and Methods Tables S3-S6, and supplemental references.

Table S1. Relative ASV abundance at family level from 16S rRNA sequencing analysis, related to Figures 1 and 3.

Table S2. Gene set enrichment analyses (GSEA) from host transcriptomics, related to Figure 4.

## Methods

### Bacterial strains and growth conditions

Bacterial strains used in this study are listed in **Table S3**. *Escherichia coli* UTSW-01 was originally isolated from feces collected from a C57BL/6 WT mouse at the UT Southwestern Medical Center. Isogenic mutants were generated using lambda red recombineering.^89,90^ For competition experiments, parental strains were chromosomally tagged with an antibiotic selection marker at the miniTN7 attachment site (*att7*) located downstream from *glmS* using lamda red recombineering.^91,92^ The plasmids and primers utilized to construct all bacterial mutants are listed in **Tables S3 and S4**. Prior to subculture for *in vitro* experiments, mouse infections, or growth curves, all bacterial strains were grown overnight in LB broth at 37°C at 250 rpm unless otherwise noted.

### Bacterial growth curves

To phenotypically assess the contribution of terminal reductases to aerobic growth, *E. coli* respiration mutants and the corresponding parental strain were grown at 37°C in LB broth under aerobic or anaerobic conditions. To assess the effects of 2-arachidonyl-glycerol (10μM or 100μM in methanol) on *E. coli* growth, bacteria were cultured in M9 minimal media supplemented with 0.2% glucose. Methanol at 1:10000 served as the vehicle control. M9 medium was prepared using M9 salts and supplemented with 2 mM MgSO_4_, 0.1 mM CaCl_2_, and 4 μg/ mL nicotinic acid. Bacterial growth was assessed by performing plate counts and via optical density readings at 600 nm using either a SpectraMax iD3 plate reader with SoftMax Pro software (version 7.1.2.) or the Thermoscientific Genesys 150 UV-Visible Spectrophotometer.

### Mice

C57BL/6 wildtype, *Cnr ^-/-^, and Il10 ^-/-^*breeding pairs were purchased from The Jackson Laboratory to generate a mouse colony at the University of South Carolina. Genotypes were confirmed with the following strain specific primer sets using LongAmp Taq 2X Master Mix (New England Biolabs): *Cnr1* common primer 5’ F: GCT GTC TCT GGT CCT CTT AAA, *Cnr1* WT 5’ R: GGT GTC ACC TCT GAA AAC AGA, *Cnr1* mutant 5’ F: AGA CTG CCT TGG GAA AAG CG; *Il10* common primer 5’ F: CTT GCA CTA CCA AAG CCA CA, *Il10* WT R: GTT ATT GTC TTC CCG GCT GT, *Il10* mutant R: CCA CAC GCG TCA CCT TAA TA. Primer sequences were obtained from The Jackson Laboratory. All mice were bred and housed in specific pathogen free conditions on a 12-hour light/dark cycle with unlimited access to water and food at the University of South Carolina. All animal protocols were approved by the University of South Carolina Institutional Animal Care and Use Committees.

### Pharmacological treatments in mouse models

To manipulate the endocannabinoid system, at the indicated time points and dosing regimens, mice were intraperitoneally (IP) injected with the Magl inhibitor JZL184 (dose range 4-40 mg/kg), the CB2 inverse agonist AM630 (dose 5 mg/kg), and/or the CB1 antagonist / inverse agonist AM281 (dose range 3-6 mg/kg) (Cayman Chemicals).^32,46,93–96^ In some experiments, mice were IP injected with WIN 55,212-2 mesylate, a CB1 / CB2 agonist (dose 5 mg/kg, Tocris).^97^ Injections were administered using a vehicle consisting of saline, DMSO, and Tween80 at an 18:1:1 ratio. In all experiments, control groups were administered the respective vehicle.

### E. coli engraftment

Enterobacteriaceae-free mice (6-11 weeks old) were engrafted with *E. coli* strain UTSW-01 via a single oral gavage with 1×10^8^ CFUs without prior antibiotic treatment. Adherent invasive *E. coli* strain NC101 was engrafted via single oral gavage with 1×10_8_ CFUs at 24 hours post treatment with streptomycin (20 mg in 100 uL of sterile PBS by oral gavage). Stable *E. coli* colonization over 3 weeks was monitored and confirmed by selective culture and targeted qPCR prior to additional treatments. At necropsy, colon tissues were collected for RNA isolation; cecal contents and feces were collected for bacterial quantification via culture, targeted qPCR, and/or 16S rRNA sequencing. *E. coli* CFUs in intestinal content homogenates were quantified by serial dilutions and plating on MacConkey agar. Mucus-associated bacteria were enumerated by quantitative selective culture from intestinal tissues as previously described.^98^

### C. rodentium infection model

Enterobacteriaceae-free mice (8-12 weeks old) were first treated via I.P. injection with JZL184 (various doses) or vehicle control. Three hours later, mice were orally infected with 1 x 10^9^ CFU of *C. rodentium* DBS100 or mock infected with sterile PBS. In experiments that included AM281 (3 mg/kg), daily treatment with this agent was initiated 24 hours prior to infection with *C. rodentium.* In experiments that included daily treatments of JZL184 and/or AM281, the agents were administered once every 24 hours until the day prior to necropsy. Mice were sacrificed for disease evaluation and cecal pathogen colonization at day 2 or 6 post infection. Fecal pellets were collected to enumerate *C. rodentium* abundance by quantitative culture on LB agar with nalidixic acid (30 *μ*g/ mL). Cecal tissues were harvested to quantify *C. rodentium* colonization, for pathology analysis, and for RNA isolation from tissues as previously described.^46^

### In vivo E. coli growth competitions

Enterobacteriaceae-free mice (6-10-weeks old) were first treated via I.P. injection with JZL184 (32 mg/kg) or vehicle control. Two hours later, mice were inoculated by oral gavage with 1×10^8^ CFUs of each *E. coli* strain at a 1:1 ratio to initiate the competitive growth experiment. *E. coli* strains were tagged with a different antibiotic resistance gene (*cat* or *kan*) to enable selective plating of each strain. Homogenized intestinal contents were serially diluted, then plated on LB agar supplemented with chloramphenicol (35 *μ*g/ mL) or kanamycin (50 *μ*g/ mL) to calculate competitive indices (CIs). CIs were calculated by dividing the CFU count of the parental strain by the CFU count of its mutant competitor.

### Gnotobiotics

Germ free WT C57BL/6 mice were maintained at the Gnotobiotic Core at University of South Carolina. Original breeder pairs were purchased from the University of Michigan Unit for Laboratory Animal Medicine Germ-Free Mouse Facility. Absence of isolator contamination was confirmed by anaerobic fecal culture onto anaerobic reducible blood agar and modified Brain Heart Infusion Agar (supplemented with 5% w/v cysteine, 1 mg/ mL vitamin K, 5 mg/ mL hemin). Fecal gDNA was extracted to confirm germ-free status using the QIAamp PowerFecal Pro DNA kit (Qiagen) followed by qPCR using the Eubacteria Universal 16S primers.

### DNA and RNA isolations

Host tissues or intestinal contents were snap frozen in liquid nitrogen at necropsy. Genomic DNA was extracted from intestinal contents using the QIAamp PowerFecal Pro DNA kit (Qiagen) per manufacturer’s instructions. Bacterial RNA was extracted from cecal contents using the RNeasy PowerFecal Pro kit (Qiagen) per manufacturer’s instructions. RNA was extracted using the TriZol method (Ambien) according to the manufacturer’s instructions. cDNA was synthesized using SuperScript III reverse transcriptase (Invitrogen) per manufacturer’s instructions.

### Targeted qPCR

qPCR was performed with a CFX Opus 384 Real-Time PCR system (Bio-Rad) using Luna Universal qPCR Master Mix (New England Biolabs). Analysis was performed using Bio-Rad CFX Manager version 3.1.1517.0823. Melting curves were assessed to ensure specificity of the PCR products. All primers used for targeted qPCR of the bacterial 16S rRNA gene are listed in **Table S5**. The relative abundance of each bacterial taxon was determined using the ΔΔ*C_T_* method normalized to the eubacteria 16S rRNA amplicon. Mammalian and *E. coli* mRNA transcripts were amplified using primers listed in **Table S6**. For mammalian gene expression, the relative abundance of each mRNA was normalized to the housekeeping genes *Gapdh* (cecum) or *Eef2* (colon). For *E .coli* gene expression, the relative abundance of each mRNA was normalized to the geometric mean of two housekeeping genes (*rpoA*, *rrs*).

### Illumina 16S rRNA sequencing

Fecal DNA isolations, library preparation, and 16S rRNA sequencing was completed at the University of Alabama Birmingham Microbiome Institutional Research Core.^99,100^ Raw sequencing data were in FASTQ format. To create amplicon sequences variants (ASVs), the Divisive Amplicon Denoising Algorithm 2 (DADA2) pipeline was used on the V4 variable region of the 16S rRNA gene and clustered at the equivalent of 100% sequence similarity.^101^ The Illumina reads were trimmed at the 240 bp and 160 bp locations for forward and reverse reads, respectively. Reads were then denoised, dereplicated, merged, and chimeric reads were removed using DADA2 standard filtering parameters. Quality control was performed to ensure no extraneous read removal during the data cleaning process. The resulting ASVs and representative reads were annotated and blasted using the Naïve Bayesian method against the SILVA database (v138.2).^102,103^ Graphical analysis and plots were created using the R packages phyloseq, ggplot2, and ggrepel.^104–106^ Alpha diversity was done based on the Shannon and Simpson indexes, which reflect both species richness and evenness within a sample. The principal coordinate analysis (PCoA) is based on Bray-Curtis beta diversity and was used to assess the between group dissimilarity in microbial diversity. This pipeline was designed to use ASVs as input and visualize the between group differences of the most abundant bacteria. Code for data analysis can be found at https://github.com/nayelilizduckworth/16S-rRNA-DADA-Analysis-Pipeline-R.

### Histopathology Analysis

Half ceca were fixed in 10% neutral buffered formalin, sectioned, and stained with hematoxylin and eosin. Inflammation was blindly assessed by measuring crypt hyperplasia using the ECHO Revolve microscope and software. The lengths of at least 5 full-length crypts per 10X field were measured in microns (6-10 fields per mouse). Representative images were taken using the ECHO Revolve microscope.

### RNA-seq and pathway analysis

Illumina library preparation and high-throughput sequencing of isolated RNA from the colon lamina propria and epithelium was completed at the Duke Sequencing and Genomic Technologies Core. Poly(A)+ libraries were generated using the KAPA mRNA Stranded HyperPrep Kit (Roche). Libraries were sequenced on a NovaSeq X Plus platform using one lane of a 25B flow cell to generate 50-bp paired-end reads. Sequencing quality control and processing were performed using nf-core/rnaseq pipeline (v3.22.2), executed with Nextflow on a high-performance computing cluster.^107,108^ Within the nf-core workflow, reads were adapter-trimmed and quality-filtered prior to alignment to the *Mus musculus* reference genome (GRCm38/mm10) using STAR. Differential expression analysis was performed in R (v4.3.2) using DESeq2 (v1.42.1).^109^ Using Wald tests, three sets of pairwise comparisons were generated, and *p-*values were adjusted by the Benjamini-Hochberg methods.^110^

Gene set enrichment analysis (GSEA)^111^ was performed using clusterProfiler (v4.10.1)^112^ and enrichplot (v1.22.0, release 20260710). For all GSEA, gene annotations were acquired from org.Mm.eg.db (3.18.0) and gene symbols were mapped to Entrez identifiers. Genes not matched with a unique Entrez ID were excluded from the ranked list and following analyses. Genes were ranked by -log10(P) signed by the direction of the log2 fold change. Enrichment was evaluated with the GSEA function using 1,000 permutations and Benjamini-Hochberg correction. Pathways with an adjusted P value (FDR) < 0.05 were considered significant. GSEA was performed individually against the following gene-set databases: KEGG (v1.42.0)^113^ and MSigDB hallmark gene sets (v25.1.0)^114^. Heatmaps were generated in Graphpad Prism.

Mitochondrial pathway enrichment was assessed against MitoCarta 3.0 MitoPathways collection for *Mus musculus.*^57^ MitoPathways organizes mitochondrial genes into a three-level hierarchy of pathways. Using org.Mm.eg.db, gene symbols were mapped to Entrez IDs. Pathways with fewer than five genes detected in the ranked list were excluded. The leading edge of each pathway was taken as the core enrichment of GSEA output, and the leading-edge fraction is defined as the leading-edge genes divided by the number of pathway genes present in the ranked list. MitoPathways pathway-level results were visualized in R using ggplot2 and ggrepel. For the OXPHOS dot plot, pathways were chosen by position within the MitoCarta hierarchy, and all Level 2 MitoPathways within the OXPHOS branch that passed the minimum gene set size threshold were included.

### 2-AG measurements and LC-MS analysis

Intestinal tissues were frozen and dehydrated by lyophilization and then weighed and homogenized in 1mL of methanol/dichloromethane (1:2, v/v) using a 5mm stainless steel bead (Qiagen) at 40 Hz for 1 hr using a TissueLyser LT (Qiagen). Homogenates were transferred to glass vials and dried under a gentle stream of N_2_ and reconstituted to a final concentration of 10 mg/mL in methanol. Lipid extractions were performed in SureStart 2mL glass vials and silicone PTFE-lined caps (Thermo Fisher Scientific). Samples were separated on a Thermo Fisher Scientific Vanquish UHPLC equipped with a Waters XBridge C18 column (2.1 x 100 mm, 3.5 µm particle size). A binary gradient of water + 0.1% formic acid (solvent A) and acetonitrile + 0.1% formic acid (solvent B) was used to perform the separation. The gradient began at 50% B and ramped to 98% B over 15 minutes, held at 98% B for 13 minutes, then re-equilibrated at 50% B for 8 minutes for a total run time of 36 minutes. All solvents were LC-MS grade and purchased from Sigma-Aldrich. Mass spectrometry was performed on a Thermo Fisher Scientific Q-Exactive HF-X quadrupole-orbitrap operating in positive mode. A reference standard of deuterated 2-arachidonoyl glycerol (D8-2-AG) (Cayman) was used to validate the identity of 2-AG in intestinal tissue extracts. The mass spectrometer performed both parallel reaction monitoring (PRM) MSMS measurements and targeted SIM measurements on the [M+H]^+^ ion of 2-AGs (*m/z* 379.2843) and D8-2-AG (*m/z* 387.3345). In PRM, the fragment ions monitored were *m/z* 287.2367 for 2-AG and a cluster of fragments at *m/z* 294.28 for D8-2-AG. 2-AG peak area was used to compare relative abundance between concentration normalized-samples.

### ATP measurements

ATP was quantified using the ATP Determination Kit (A22066, Invitrogen), a firefly luciferase-based bioluminescence assay. The kit was used according to manufacturer’s instructions. The kit was used to measure ATP in supernatant from homogenized colon mucosal tissue and individual sample luminescence was measured on the SpectraMax iD3 plate reader. ATP values were normalized to tissue weight, and data are reported as absolute or relative ATP fold-change versus the mean of the control group.

### Statistical analysis

*P-*values were calculated using Mann-Whitney or Student’s *t* test when two experimental groups were compared, one-way ANOVA or Kruskal-Wallis when three or more experimental groups were compared, and two-way ANOVA when two independent variables were included. CFU counts generated from bacteria culture and competitive indices were routinely log transformed for normalization. Unless otherwise noted, data are represented as mean ± SEM. Graphs were generated and statistics were performed using GraphPad Prism version 10.6.0 or R Version 2025.09.0+387.

## Notes

### Competing Interest Statement

The authors have declared no competing interest.

