## Supplemental Materials for "Cannabinoid receptor 1 activity promotes microbial dysbiosis by stimulating the oxygen-dependent expansion of Enterobacteriaceae"

### Supplemental Figures

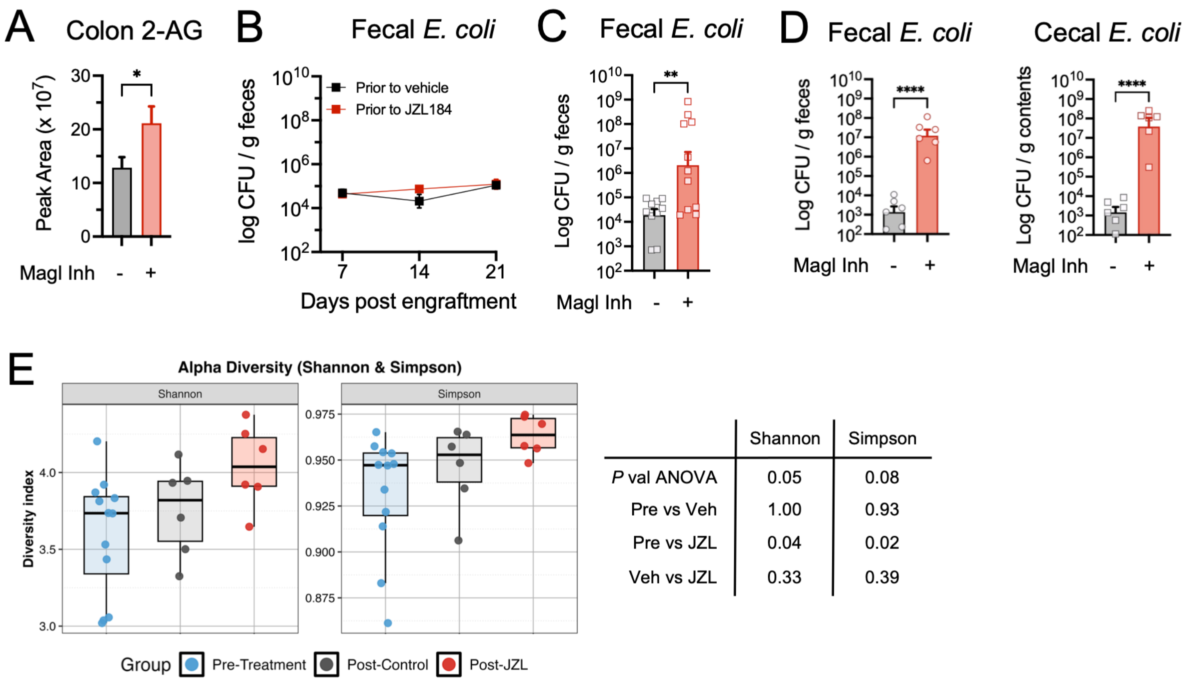


| | Shannon | Simpson |
|---|---|---|
| *P* val ANOVA | 0.05 | 0.08 |
| Pre vs Veh | 1.00 | 0.93 |
| Pre vs JZL | 0.04 | 0.02 |
| Veh vs JZL | 0.33 | 0.39 |

**Fig. S1. Related to Fig. 1.**

A) Mice were acutely treated with vehicle or Magl inhibitor (32 mg/kg) ($n$ = 6-7 per group). Detection and quantification of 2-AG by LC-MS in colon tissues at 15-17 hours post treatment.

B) Longitudinal fecal *E. coli* culture following engraftment and prior to the vehicle or Magl inhibitor treatment. Corresponds with data presented in Fig 1D-G.

C) Male mice engrafted with *E. coli* were treated with vehicle or Magl inhibitor at 32 mg/kg ($n$ = 10-11 per group). Data depict fecal *E. coli* CFUs at 12 hours post treatment. Student's t test.

D) Post treatment *E. coli* CFUs in feces and cecal contents. Corresponds with data presented in Fig 1D-G. Student t-test.

E) Alpha diversity metrics of 16S rRNA sequencing data presented in Fig 1D-G. Lines are on the median. One-way ANOVA with Bonferroni multiple comparison test.

* $p < 0.05$, ** $p < 0.01$, **** $p < 0.0001$

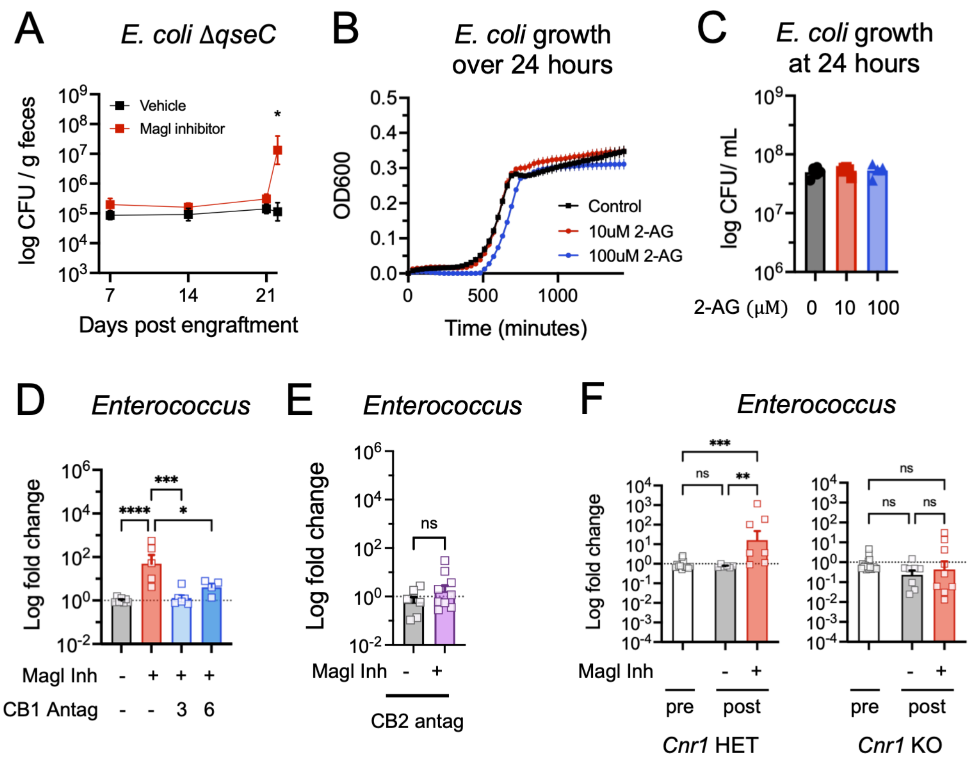


**Fig. S2. Related to Fig. 2.**

A) Female mice were engrafted with an *E. coli qseC*-deficient mutant, then treated with vehicle or Magl inhibitor (32 mg/kg) ($n$ = 6-9 per group). Fecal *E. coli* CFUs post initial engraftment. Two-way ANOVA with Sidak's multiple comparison test.

B) Growth curves and C) CFU counts at 24 hours of *E. coli* UTSW-01 in the presence of increasing concentrations of 2-AG in M9 minimal medium with glucose as the sole carbon source.

D-F) Post treatment relative abundance of fecal *Enterococcus*, as determined by targeted qPCR, with D) CB1 antagonist (3 or 6 mg/kg) in combination with Magl inhibitor (32 mg/kg) in WT mice, E) CB2 antagonist (5 mg/kg) alone or in combination with Magl inhibitor (32 mg/kg) in WT mice, or F) Magl inhibitor (32 mg/kg) in *Cnr1* HET or KO mice. One-way ANOVA with Tukey's multiple comparison test.

* $p < 0.05$, ** $p < 0.01$, *** $p < 0.001$, **** $p < 0.0001$

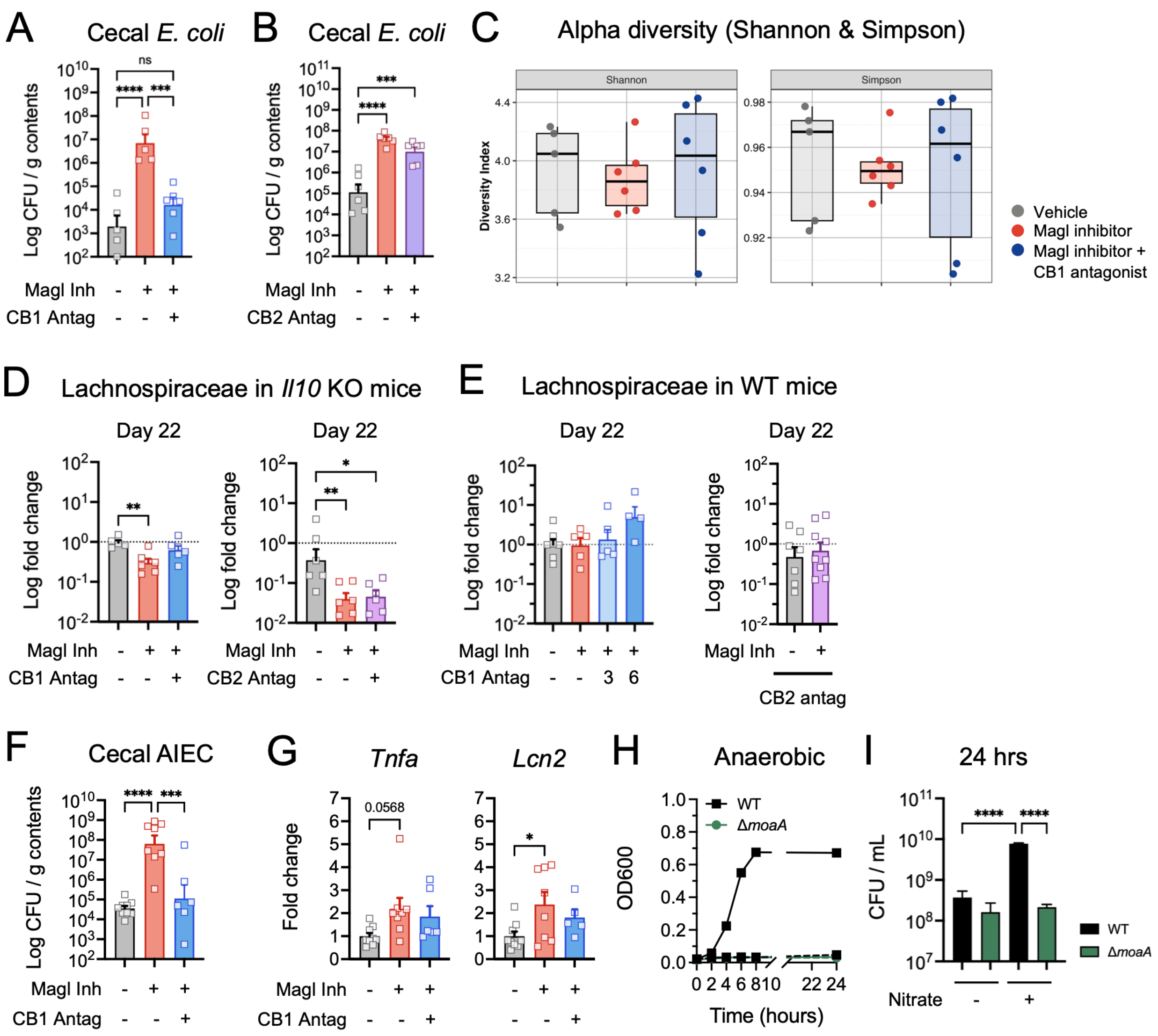


**Fig. S3. Related to Fig. 3.**

A-E) Female *Il10* KO mice (n = 5-6 per group) were engrafted with *E. coli,* then three weeks later were treated with vehicle, Magl inhibitor (32 mg/kg), or CB1 antagonist (3 mg/kg) or CB2 antagonist (5 mg/kg) in combination with Magl inhibitor.

A-B) Post treatment *E. coli* CFUs in cecal contents at day 22. Corresponds with data presented in Fig 3A. One-way ANOVA with Tukey's multiple comparisons test.

C) Alpha diversity metrics of 16S rRNA sequencing data presented in Fig 3B-C.

D) Post treatment relative abundance of fecal Lachnospiraceae relative to vehicle group, as determined by targeted qPCR, in *Il10* KO mice*.* One-way ANOVA with Tukey's multiple comparisons test.

E) Post treatment relative abundance of fecal Lachnospiraceae relative to vehicle group, as determined by targeted qPCR, in WT mice. The numbers indicate the dose of CB1 antagonist in mg/kg. One-way ANOVA with Tukey's multiple comparisons test (left). Student t test (right).

F-G) Female *Il10* KO mice (n = 6-9 per group) were engrafted with AIEC*,* then three weeks later were treated with vehicle, Magl inhibitor (32 mg/kg), or CB1 antagonist (6 mg/kg) in combination with Magl inhibitor. F) Post treatment AIEC CFUs in cecal contents. G) Host expression of *Tnfa* and *Lcn2* in colon tissues relative to vehicle. Corresponds with data presented in Fig 3D-F. One-way ANOVA with Tukey's multiple comparisons test.

H) Anaerobic growth curves and I) CFU counts at 24 hours of *E. coli* strains in the presence (solid lines) or absence (dashed lines) of nitrate. One-way ANOVA with Tukey's multiple comparisons test.

* $p < 0.05$, ** $p < 0.01$, *** $p < 0.001$, **** $p < 0.0001$.

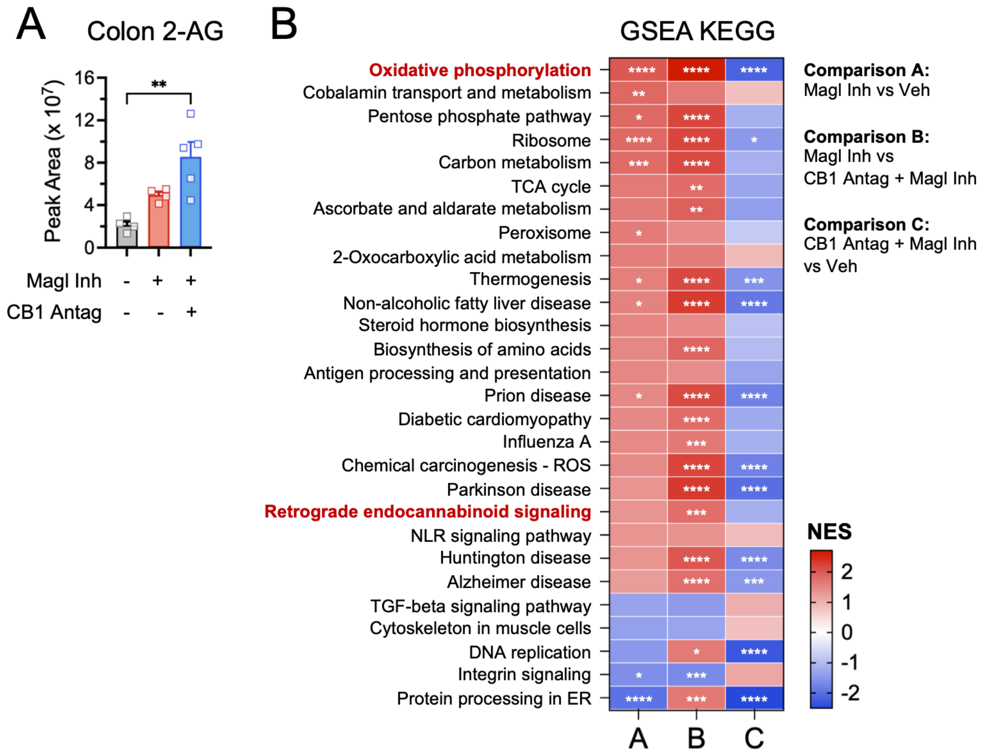


**Fig. S4. Related to Fig. 4.**

A) Detection and quantification of 2-AG by LC-MS in colon tissues at 3 hours post-treatment with vehicle, Magl inhibitor (32 mg/kg), or CB1 antagonist (6 mg/kg) in combination with the Magl inhibitor. Related to Fig 6A-D. Kruskal-Wallis with Dunn's multiple comparisons test.

B) Gene set enrichment analysis (GSEA) using the GSEA gene set collection. Only enriched pathways with FDR < 0.2 in comparisons A and B were included. * FDR < 0.05, ** FDR < 0.01, *** FDR < 0.001, **** FDR < 0.0001

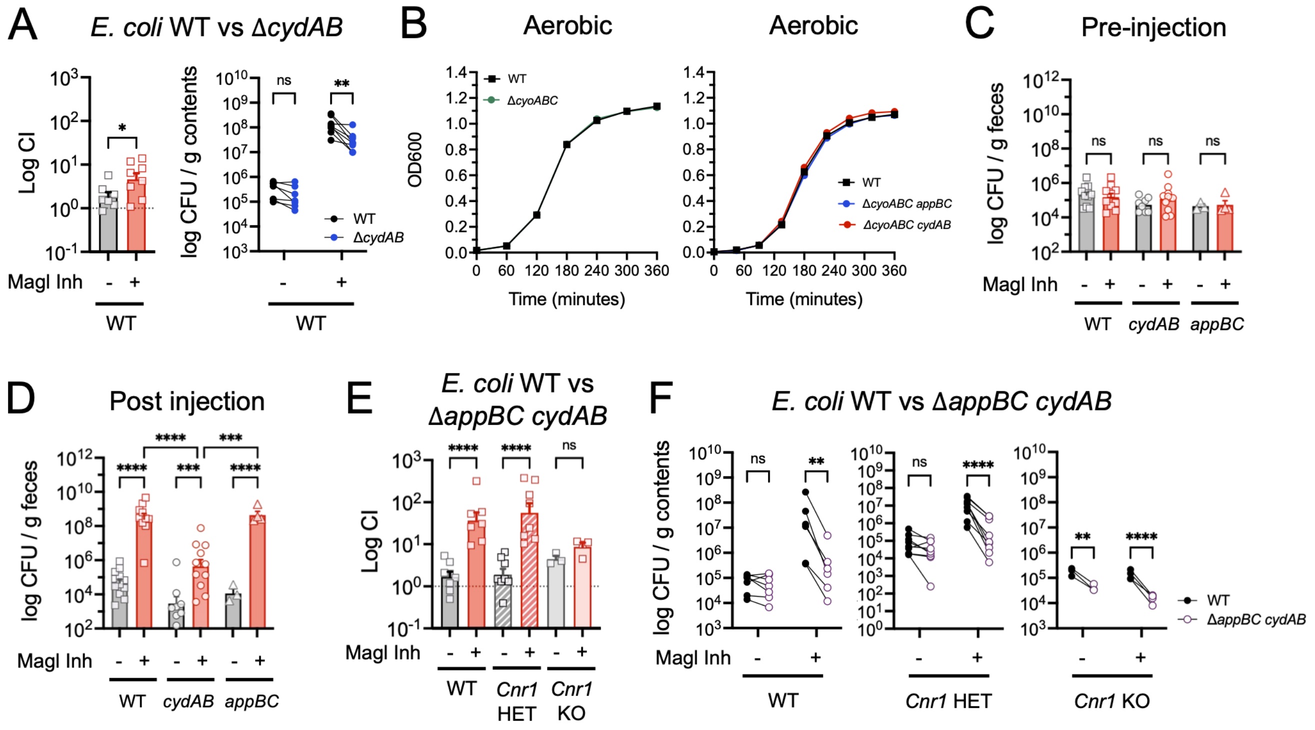


**Fig. S5. Related to Fig. 6.**

A) Female mice (n = 7-8 per group) were treated with a single dose of vehicle or Magl inhibitor (32 mg/kg), then gavaged 2 hours later at a 1:1 ratio with the indicated *E. coli* isogenic mutant and parental strain. Competitive indices (parental strain vs mutant) were calculated from *E. coli* CFUs cultured from cecal contents. Student's t test (first panel); Two-way ANOVA with Sidak's multiple comparison test (second panel). Corresponds with data presented in Fig 6C.

B) Growth curves of individual *E. coli* strains under aerobic conditions.

C-D) Male and female mice (*n* = 3-11 per group) were engrafted with the indicated *E. coli* isogenic mutant or parental strain, then three weeks later were treated with a single dose of vehicle or Magl inhibitor (32 mg/kg). D) Pre-treatment and E) post-treatment *E. coli* CFUs in feces. Two-way ANOVA with Sidak's multiple comparison test.

E-F) Male and female WT, *Cnr1* HET, or *Cnr1* KO mice (n = 3-8 per group) were treated with vehicle or Magl inhibitor (32 mg/kg), then gavaged 2 hours later at a 1:1 ratio with the indicated *E. coli* isogenic mutant and parental strain. E) Competitive indices (parental strain vs mutant) were calculated from F) *E. coli* CFUs cultured from cecal contents. Corresponds with data presented in Fig 6F-G. Two-way ANOVA with Sidak's multiple comparison test.

* $p < 0.05$, ** $p < 0.01$, *** $p < 0.001$, **** $p < 0.0001$

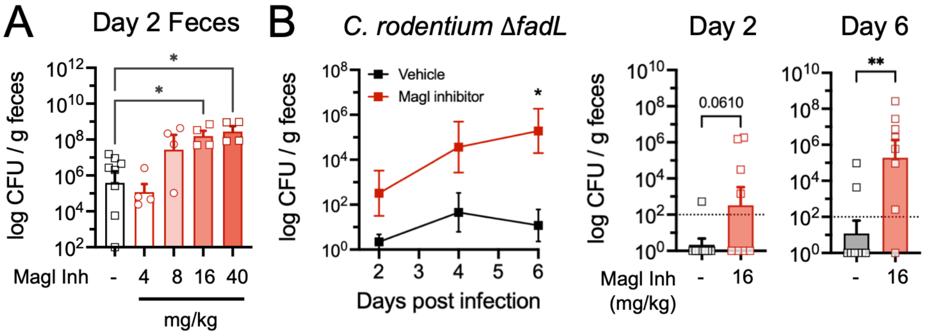


**Fig. S6. Related to Fig. 7.**

A) Female mice (*n* = 4-8 per group, pooled from 2 cohorts) were infected with *C. rodentium* and administered daily injections of vehicle or Magl inhibitor. *C. rodentium* CFUs in feces at day 2 post infection. Corresponds with data presented in Fig 3B. One-way ANOVA with Tukey's multiple comparisons tests relative to vehicle.

B) Female mice (*n* = 8 per group, pooled from 2 cohorts) were gavaged with *C. rodentium ΔfadL* and administered daily injections of vehicle or Magl inhibitor (16 mg/kg). B) Fecal *C. rodentium ΔfadL* CFUs. Two-way ANOVA with Sidak's multiple comparison test. *C. rodentium ΔfadL* CFUs in feces at days 2 and 6 post infection. Dotted line represents limit of detection. Student t-test.

* $p < 0.05$, ** $p < 0.01$

### Supplemental Materials and Methods

#### Table S3. Bacterial strains and plasmids

| **Strain Name** | **Species** | **Reference** |
|---|---|---|
| UTSW-01 | *Escherichia coli* | This study. |
| NC101 | *Escherichia coli* | Kim SC 2005[1] |
| NC101 Δ*moaA* | *Escherichia coli* | This study. |
| UTSW-01 Δ*cydAB* | *Escherichia coli* | This study. |
| UTSW-01 *cydAB*::*kan* | *Escherichia coli* | This study. |
| UTSW-01 Δ*appBC* | *Escherichia coli* | This study. |
| UTSW-01 *appBC*::*kan* | *Escherichia coli* | This study. |
| UTSW-01 Δ*cyoABC* | *Escherichia coli* | This study. |
| UTSW-01 *cyoABC*::*kan* | *Escherichia coli* | This study. |
| UTSW-01 Δ*appBC cydAB::cm* | *Escherichia coli* | This study. |
| UTSW-01 Δ*cyoAB cydAB::cm* | *Escherichia coli* | This study. |
| UTSW-01 Δ*cyoABC appBC::cm* | *Escherichia coli* | This study. |
| UTSW-01 *qseC*::*kan* | *Escherichia coli* | This study. |
| UTSW-01 *att7*::*kan* | *Escherichia coli* | This study. |
| UTSW-01 *att7*::*cm* | *Escherichia coli* | This study. |
| DBS100 | *Citrobacter rodentium* | Ellermann M 2020[2] |
| DBS100 Δ*fadL* | *Citrobacter rodentium* | Ellermann M 2020[2] |
| **Plasmid Name** | **Purpose** | **Reference** |
| pKD3 | For lambda red recombineering | Datsenko KA 2000[3] |
| pKD4 | For lambda red recombineering | Datsenko KA 2000[3] |
| pKD46 | For lambda red recombineering | Datsenko KA 2000[3] |
| pCP20 | For lambda red recombineering | Datsenko KA 2000[3] |

### Table S4. Primers used for construction of bacterial mutants

| Purpose | Sequence (5' → 3') |
|---|---|
| *cydAB* KO primers | **F:**TGCGTGTGACCCTTTCTAACGCGGTTCACTCTCGGAGTCTTCATGCGATGAGCAAGGAGTCATGATGTTAGTGTAGGCTGGAGCTGCTTC<br>**R:**CTTAGTACAGAGAGTGGGTGTTACGTTCAATATCTTCTTTGGTGATACGACCGAACATTTTCCAGTAACATATGAATATCCTCCTTAGTT |
| *cydAB* confirmation primers | **F:** GCGTGTGACCCTTTCTAACGG<br>**R:** ATCAGATGTCTTCTTGACCGG |
| *qseC* KO primers | **F:**GGTGAGAAATGAAATTTACCCAACGTCTTAGTCTGCGAGTCAGGCTGACGCTAATCTTTTTAATTCTGGTGTAGGCTGGAGCTGCTTC<br>**R:**TATGAGAGTCGTTTTAACGGCTCTCATAGACAGAGAAGTTACCAGCTTACCTTCGCCTCAAATCCACCCATATGAATATCCTCCTTAGTT |
| *qseC* confirmation primers | **F:** GTATACCTGGGACGAAGAGG<br>**R:** GCGGGAAATGTTCGAGTTTTGG |
| *moaA* KO primers | **F:**GTAAAGATTGCCACCAGGGCGAAGGAAGAAATGACTTCGCCTCCCGTATCTGGAAAGGTGTACATGGCTGTGTAGGCTGGAGCTGCTTC<br>**R:**TGACGTTTTAGCCGCCAATGTACGATAAGTTTTGCGTAATACCGGTGTTGTTTTGATGCAGGAAATGCATATGAATATCCTCCTTAGTT |
| *moaA* confirmation primers | **F:** CTCTAGCCTCTGCACCTGGG<br>**R:** ATCGCGCAGATAGTGACCGG |
| *appBC* KO primers | **F:**CAGGATAAAGAGGGAGATCTACCATTATTGGGTTATTTTTCTCTCTTCGCCTACAGGAGTGCGCATGTGGGTGTAGGCTGGAGCTGCTTC<br>**R:**AACAAAATGCCGACGAACCAAAGTAAATACCACATTGTTCCTGCTCCTTAGTACAGCTCGTTTTCGTTCATATGAATATCCTCCTTAGTT |
| *appBC* confirmation primers | **F:** GCAATCGGCGAATACCTCTGG<br>**R:** GCCATACCAACACAAGGGTGG |
| *cyoABC* KO primers | **F:**TTAAACGCCACCAGATCCCGTGGAATTGAGGTCGTTAAATGAGACTCAGGGTGTAGGCTGGAGCTGCTTC<br>**R:**TGCGTTAGAATGACTCATTACATCGCCCCCATCAGATAAACAACAGTGAACATATGAATATCCTCCTTAGTT |
| *cyoABC* confirmation primers | **F:**TGTTTACAGTAATGTAACCTTCC<br>**R:** CCGTCAGGATTATCGACAGG |
| Antibiotic cassette insertion into the *att7* site | **F:**GAATGTCTTCAGTCCGATTTAAATAAGAGTTTATATTCAGTTAATTACAAATATTAATAACGAAGAGATGGTGTAGGCTGGAGCTGCTTC<br>**R:**ATGTTTTTAATCAAACATCCTGCCAACTCCATGTGACAAACCGTCATCTTCGGCTACTTTTTCTCTGTCACATATGAATATCCTCCTTAGTT |
| Antibiotic tagging confirmation primers | **F:** AATCAAACATCCTGCCAACTCC<br>**R:** GAAAGCGCTCATCGATAAGG |

#### Table S5. Bacterial qPCR primers

| Target | Sequence (5' → 3') | Citation |
|---|---|---|
| Eubacteria (Universal 16S) | **F:** TGTCGTCAGCTCGTGTCGTGA<br>**R:** ATCCCCACCTTCCTCCGGT | This study |
| Enterobacteriaceae 16S rRNA gene | **F:** GTGCCAGCMGCCGCGGTAA<br>**R:** GCCTCAAGGGCACAACCTCCAAG | Zhu W 2018[4] |
| *Enterococcus spp.* 16S rRNA gene | **F:** AGAAATTCCAAACGAACTTG<br>**R:** CAGTGCTCTACCTCCATCATT | Ryu H 2013[5] |
| Lachnospiraceae 16S rRNA gene | **F:** GTAAAGCTCTATCAGCAGGGAAGA<br>**R:** CGACAGTTTCCAATGCAGTACCGG | FP: Chen L 2017[6]<br>RP: This study. |

#### Table S6. Mammalian and *E. coli* qPCR primers

| Target Gene | Sequence (5' → 3') | Citation |
|---|---|---|
| *Gapdh* | **F:** GGTGAAGGTCGGAGTCAACGGA<br>**R:** GAGGGATCTCGCTCCTGGAAGA | Ellermann M 2020[2] |
| *Eef2* | **F:** TGTCAGTCATCGCCCATGTG<br>**R:** CATCCTTGCGAGTGTCAGTGA | Eissa N 2016, Scientific Reports |
| *Nos2* | **F:** TTGGGTCTTGTTCACTCCACGG<br>**R:** CCTCTTTCAGGTCACTTTGGTAGG | Ellermann M 2020[2] |
| *Tnfa* | **F:** AGCCAGGAGGGAGAACAGAAAC<br>**R:** CCAGTGAGTGAAAGGGACAGAACC | Hughes E 2017, CHM |
| *Lcn2* | **F:** AAGGCAGCTTTACGATGTACAGC<br>**R:** CTTGCACATTGTAGCTCTGTACC | Chassaing B 2012, PLoS One |
| *rpoA* | **F:** GTGACCCTTGAGCCTTTAGAG<br>**R:** ACACCATCAATCTCAACCTCG | Ellermann M 2020[2] |
| *rrs* | **F:** GTTAATACCTTTGCTCATTGA<br>**R:** ACCAGGGTATCTAATCCTGTT | Ellermann M 2020[2] |
| *cydA* | **F:** GTTCTGGGTGATGAATCCGG<br>**R:** TCTTCCTGATCAGGAATGCC | This study |
| *cyoA* | **F:** TAAGTACCGGGTTCGTTGG<br>**R:** AAGTGGTTTCCATGGACTGG | This study |
| *appB* | **F:** TGGGAAGGTAATCAGGTCTGG<br>**R:** GATGACCAGTCCGGCATCC | This study |
